# Identifying Selectivity Filters in Protein Biosensor for Ligand Screening

**DOI:** 10.1101/2023.07.11.548514

**Authors:** Mohammad Sahil, Jayanti Singh, Subhankar Sahu, Sushant Pal, Ajit Yadav, Ruchi Anand, Jagannath Mondal

## Abstract

Specialized sensing mechanisms in bacteria enable the identification of cognate ligands with remarkable selectivity in highly xenobiotic-polluted environments, where these ligands are utilized as energy sources. Here, via an integrated all-atom computer simulation, biochemical assay and isothermal titration calorimetry approaches we determine the molecular basis of MopR, a phenol biosensor’s complex selection process of ligand entry. Our results reveal a set of strategically placed selectivity filters along the ligand entry pathway of MopR. These filters act as checkpoints, screening diverse aromatic ligands at the protein surface based on their chemical features and sizes. Ligands meeting specific criteria are allowed to enter the sensing site in an orientation-dependent manner. Sequence and structural analyses demonstrate the conservation of this ligand entry mechanism across the sensor class, with individual amino acids along the selectivity filter path playing a critical role in ligand selection. Together, this investigation highlights the importance of interactions with the ligand entry pathway, in addition to interactions within the binding pocket, for achieving ligand selectivity in biological sensing. The findings enhance our understanding of ligand selectivity in bacterial phenol biosensors and provide insights for the rational expansion of the biosensor repertoire, particularly for the biotechnologically relevant class of aromatic pollutants.

## Introduction

Bacterial transcription initiation mainly occurs via two diverse RNA polymerases, namely σ70 and σ54. While σ70 polymerase transcribes housekeeping genes and does not require any external activation to form transcriptionally competent open complex, the alternate polymerase σ54 require regulatory proteins, typically AAA+ ATPases, that aid in converting the closed RNA polymerase complex to an active open state.^1,2^ External stimuli and environmental cues trigger σ54 RNA polymerases that then govern several cellular processes ranging from specific transport systems, alternative carbon catabolism as energy source, production of extracellular structures, and virulence determinants.^3,4^ The AAA+ sub-class of σ54 activators proteins, also known as enhancer binding proteins (EBPs), possess a modular architecture and it is via their ubiquitous central AAA+ ATPase domain that they assemble into competent ATPase motors that in turn activate the σ54 RNA polymerase holoenzyme. ^5,6^ The assembly of the ATPase domain however, is regulated by the N-terminal signal sensing domain of EBPs.^7,8^ This N-terminal domain is a specialized domain that controls the whole downstream relay via sensing appropriate external stimuli. They also harbor a C-terminal DNA binding domain that binds to a specific upstream binding sequence located ∼200 bases upstream of the σ54 polymerase assembly complex enables them to come in close proximity of the RNA polymerase assembly. ^1,9^

While, the central AAA+ and the C-terminal DNA binding domains are mostly conserved it is the N-terminal domain that interfaces with the environment and determine the type of process the σ54 assembly activates.^8,10^ The N-terminal domain functions as environmental sensors, hence also called as sensor domain. In this regard organisms such *Pseudomonas* and *Acinetobacter* that survive in harsh environmental conditions and compete for nutrients have devised sophisticated sensors for pollutants such as benzene, toluene, phenol, xylenols and bi-phenyls etc.^11,12^ The sensor domains of these proteins are designed to sense these pollutants which then can be catabolized as an alternate energy source.^10,13,14^ Exploiting these sensory proteins several groups have designed protein or synthetic biology based whole cell biosensors as well as engineered these sensor proteins to devise readouts for a plethora of hydrocarbons.^15–17^ A striking feature of the natural sensory proteins is that they are very specific for the molecule they sense. For instance, a phenol sensing domain does not accept a similar aromatic such as benzene or another phenolic compounds such as dimethylphenols.^13,18^ Thus, to effectively cover the chemical space such that this sensor scaffold can be exploited to engineer a wide array of sensors the problem of ligand selection is an important one.

Although the nature’s algorithm of protein-ligand complementarity is efficient, it requires the ligand to first reach the binding pocket, where ligand selection takes place after binding.^19^ This in turn influences the residence time of ligand which defines the preference for different ligands.^20,21^ Therefore, a question that arises is: how does enzyme allow entry of their cognate ligands, especially when the binding pocket is deeply buried as in sensor domains of AAA+ class? Are there multiple selectivity filters that preclude binding/entry of random ligands and allow for a transient path that facilitates passage of only the cognate ligands or the protein opens up to directly accommodate the ligand into the pocket. Several powerful algorithms have been proposed to address this protein-ligand recognition problem where the geometrical constraints along with protein dynamics are used to identify the optimal ligand entry path as well as final binding pose of ligand.^22^ The problem is simpler in systems such as Hsp70, Maltose Binding Protein (MBP) and adenylate kinase (AdK) where explicit pathways are already visible and additional gating filters could be identified to allow for entry of the correct ligand.^23–25^ However, in majority of the cases, it is less obvious how protein opens up and selects the ligand of choice, hence a significant thrust in understanding the ligand selection problem is imperative.^26,27^

In this work we have employed MopR from *Acinetobacter calcoaceticus* as a model system (Figure 1), a very important malleable system that is being developed for environmental pollution based biotechnological applications, to shed light on the ligand selection problem.^15–17^ MopR is a natural phenol sensor and recent X-ray structure of the sensor domain shows that it is a dimer with two buried phenol binding sites.^13^ Previous ITC studies have helped generate a reliable database to understand it’s substrate scope. ^15^ However, as entailed earlier, how the ligands reach a buried pocket and the mechanism of rejection or acceptance of substrates remain unclear. Here, we carry out conformational dynamics undertaking identification and quantification of the entire binding process and binding-competent encounter complexes and intermediate states. Long unbiased Molecular Dynamics (MD) simulations that unraveled the path chartered by small molecule phenol to reach from bulk to sensor pocket of MopR in real time were performed. Insights developed from mechanistic understanding of the binding process were further substantiated by experiment. Insights from this study purported us to propose a general model for aromatic pollutant sensing mechanism which can guide future works in fine tuning the selectivity and sensitivity of the aromatic sensor class, an important arena considering the grave problem of environmental pollutant sensing. This work also sheds lights on the general mechanism adopted by proteins to allow selective entry of substrates/ligand to deeply buried active sites.

**Figure 1:**
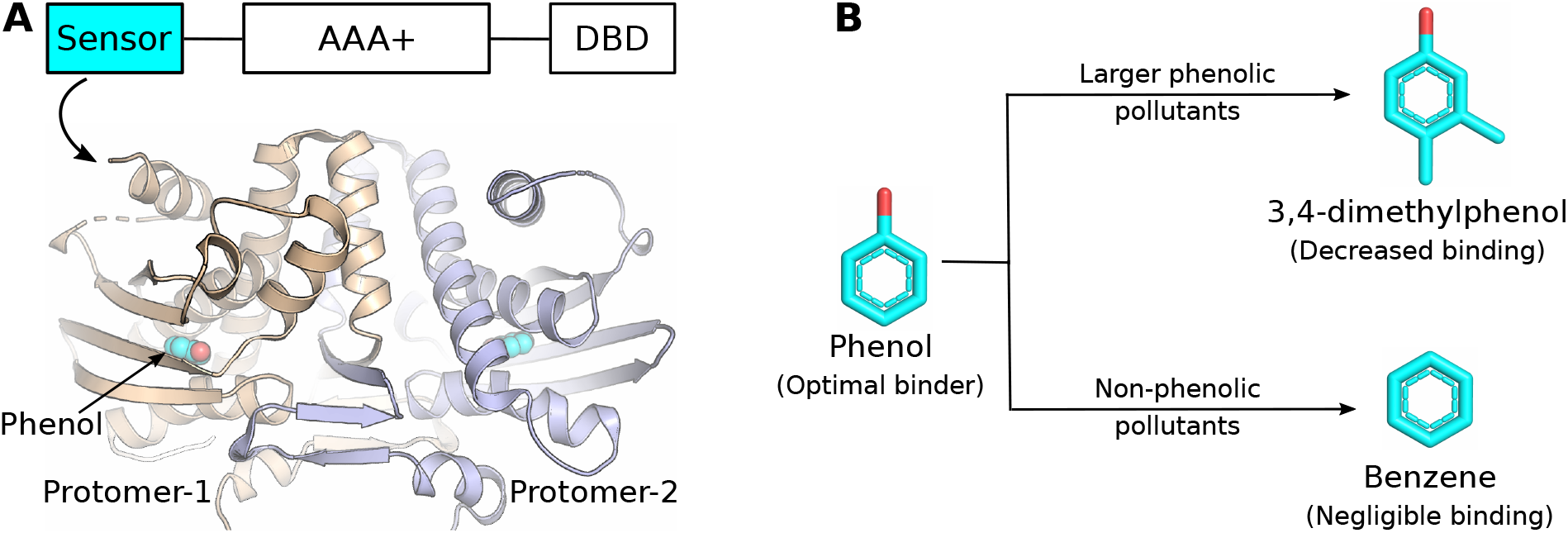
The selective MopR. (A) Homodimeric N-terminal sensor domain (residues 1-229) of phenol sensor MopR corresponding to pdb id 5KBE. Schematic on top represents the three domains of full length MopR. (B) Ligand selection profile of MopR showing size-based and phenol-based ligand selection.

## Results

### Long unbiased simulations capture the phenol binding to MopR at Real time

In a bid to elucidate the key determinants of the selective sensing ability by MopR’s sensor domain, our investigation first focused on discovering the possible binding pathway(s) of phenol in the solvent-inaccessible cavity of MopR. Towards this end, we spawned a series of all-atom unbiased MD trajectories to simulate the event of phenol diffusing around dimeric MopR in solvent. In particular, 12 binding trajectories were initiated with a set of randomly placed phenol molecules at experimentally prescribed^13^ protein/ligand ratio of 1:5. Both visual inspection and the time profile of root-mean-squared deviation (RMSD) of the simulated pose of phenol (Figure 2A) in reference with the crystallographic bound pose (PDB ID: 5KBE) helped in tracking the location of the ligand relative to the native pose. As an encouraging development, we found that, in 11 out of 12 simulations, at least one of the diffusing phenol molecules successfully identified one of the native binding pocket of MopR and remained bound for rest of the simulation period. In particular, in two instances, the buried cavity of both the protomers of dimeric MopR got phenol-bound, thereby totalling 13 binding events (labelled as Traj 1-13). The duration of simulated binding events ranged between 1.2 to 4 *µ*s. As depicted by RMSD profiles of the ligand (two representative trajectories in Figure 2A and others in Figure S1-2), the eventual simulated phenol-bound pose converged within angstrom-level precision with the crystallographic pose (Figure 2B). The results indicate that the simulations have been able to capture the binding process of phenol-sensing event by MopR at atomistically precise spatial and temporal resolution in real time. Availability of these high resolution trajectories allowed us to visualize and hypothesize the most likely pathways that the ligand had utilised in tracing the native binding pocket of MopR.

**Figure 2:**
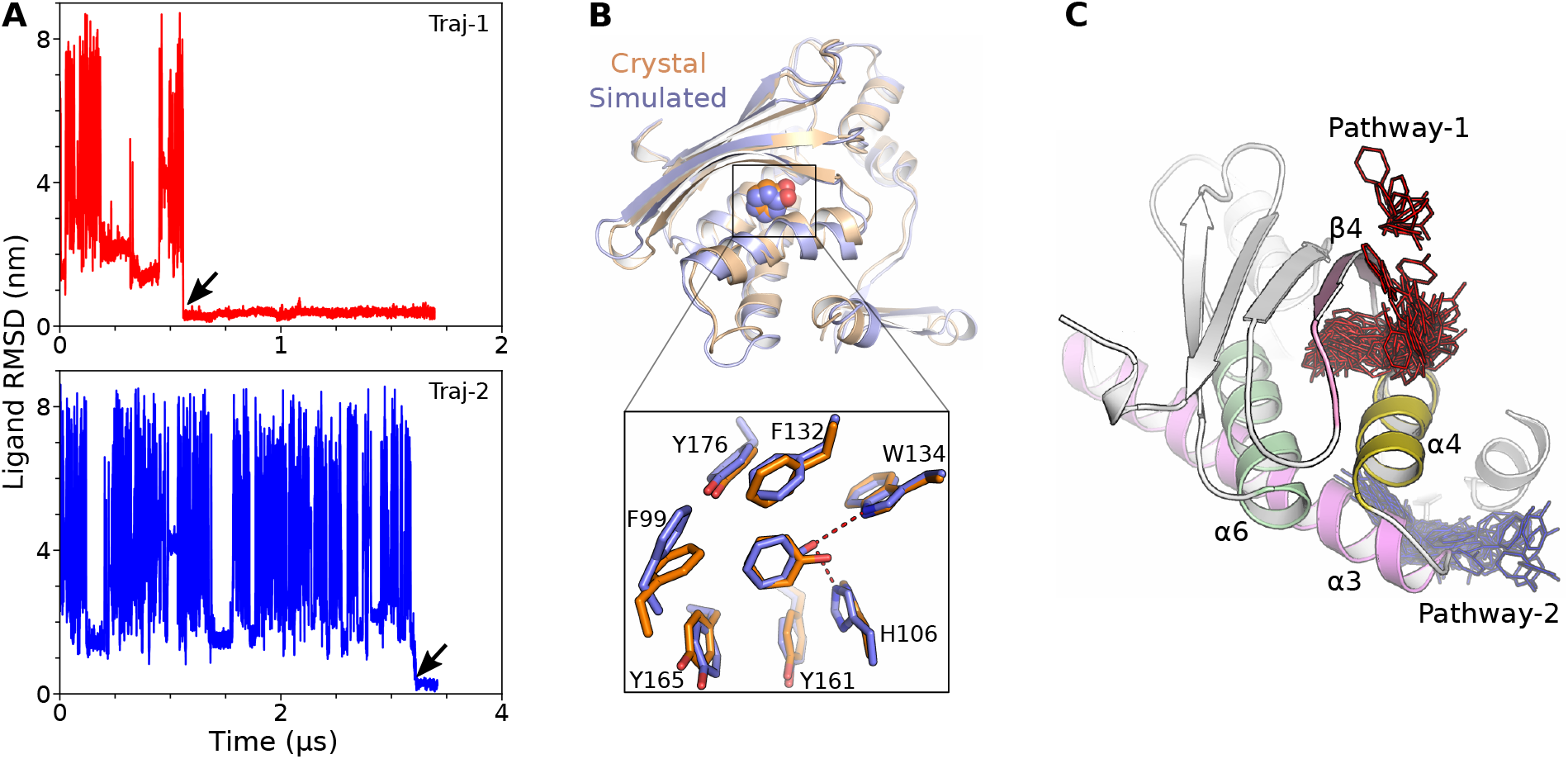
Binding pathways. (A) RMSD time profile of two representative binding trajectories labelled as Traj1,2. Arrow indicate binding. (B) Overlay of bound conformations of MD derived pose and crystal structure. Inset represent the overlay of bound state and binding pocket. (C) Two binding pathways in MopR as observed in unbiased binding simulations wherein the pathway-1 is major pathway. Only one of the protomer of dimeric MopR is shown for clarity. (dimer in Figure S3)

The simulation trajectories revealed that majority of the binding events (12 out of 13 binding events) occurred via a single dominant pathway (Figure 2C, referred here-in as ‘Pathway-1’, Movie1) which involved phenol insertion into buried binding pocket in-between *α*4 (4*^th^* helix) and *β*4 (4*^th^* strand). The other remaining binding event (one out of 13) adopted a different pathway (Figure 2C, referred here-in as ‘Pathway-2’, Movie2), which involved ligand entry to the binding pocket in between *α*4 and *α*3-6 helices. The average ligand binding time through pathway-1 was found to be around 2 times faster (within 1.5 *µ*s) than pathway-2 (3.2 *µ*s); together giving an estimated average on-rate constant (*k_on_*) of 3.5 *×* 10^7^*M ^−^*^1^*s^−^*^1^ (Figure S1), at par with reported *k_on_* values for ligand binding in buried pocket for other systems.^26,27^ The identification of aforementioned major dominant ligandbinding pathway prompted us to delve deeper into the molecular mechanism underlying the dynamics of recognition event and its potential ligand-selectivity.

### Simulations and experiments identify multiple residue barriers to buried cavity controlling ligand binding

Enroute to buried binding pocket through major pathway-1, the ligand encounter two major barriers imposed by residues of *α*4, *β*4 and binding pocket itself. Firstly, the ligand approaches the hydrophobic surface between *α*4 and *β*4, in particular at residues M100, V116 and L119, in a specific orientation such that aromatic moiety of ligand interacts with hydrophobic protein surface (Figure 3, S3). We surmised that ligand entry would require an opening in the forming of an ‘encounter-complex’. The simulation trajectories identified two possible mechanisms leading to formation of an encounter complex: one set of trajectories involved a ligand induced pocket gate opening, while in rest of the trajectories, the ligand entry was facilitated by spontaneous creation of a transient channel.

**Figure 3:**
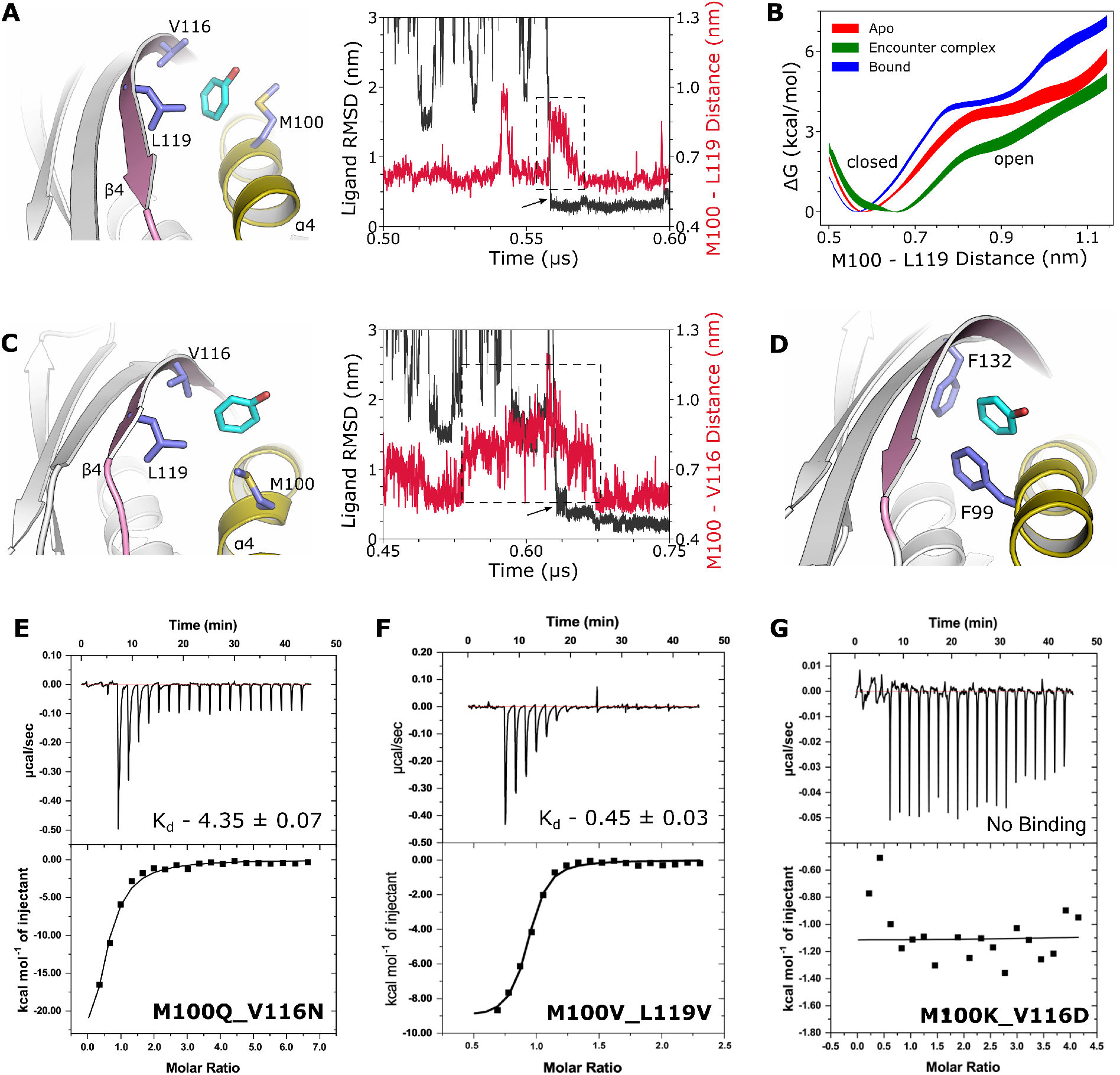
Binding mechanism. (A) Ligand entry into binding pocket via M100-L119 pocket-gate opening. Inset represent the opening while arrow indicate ligand entry. Opening at ∼0.55 represent unsuccessful binding attempt by different ligand. (B) Free energetics of pocket-gate opening in different states of ligand. Width of curve represent the error bar. (C) Ligand entry via M100-V116 channel opening. (D) Ligand encounter of F99 and F132 barrier. (E-G) The ITC binding data for different pocket gate mutants of MopR. All the *K_d_* values are represented in *µ*M. Introduction of hydrogen bonding (E) and or salt bridge (G) progressively decrease and abrogates binding respectively.

For half of the trajectories, ligand induced the opening of M100-L119 pocket-gate, which involved the outward movement of M100 sidechain away from L119 allowing ligand entry (Figure 3A). A good correlation between the increase in M100-L119 distance profile and concurrent decrease in ligand-RMSD at around same time-point in these trajectories suggested that the presence of ligand is crucial for such gate opening. For a quantitative characterization, the free-energetic requirement for induced pocket-gate opening was measured along center-of-mass (COM) distance between side chains of M100 and L119 (Figure 3B). It was found that pocket-gate opening is separated by a free-energetic barrier and the free-energy requirement for closed (0.6) to open (0.9 nm) transition are 3.77 *±* 0.21 and 2.59 *±* 0.23 kcal/mol for ‘apo’ and ‘encounter-complex’ respectively. The ligand presence lowers the free energy requirement, hence we concluded that the ligand helps in inducing the pocket gate opening. Additionally, once the ligand gets bound in the native pocket, pocket-gate opening requires more energy and gets tightly closed fulfilling the necessary characteristic of ‘high sensitivity’ for MopR as a potential candidate for phenol biosensor.

In second half of the trajectories representing major pathway-1, M100 spontaneously moves away from V116 (not L119) creating a transient channel between *α*4 and *β*4, paving the way for ligand entry to the binding pocket (Figure 3C). As shown by the time profile of M100-V116 distance as well as ligand RMSD, this transient channel opened spontaneously and independently of ligand entry. We believe the intrinsic flexibility of M100 due to presence of kink in *α*4 (as it is constituted of the residues having lowest helix propensities like G102, P103) contributes to this transient yet spontaneous gate opening. ^28,29^

After surpassing the initial obstacle set by M100, V116, and L119, the path of the ligand towards the binding pocket beyond the encounter complex was additionally constrained by a second barrier established by F99 and F132 (Figure 3D). These residues can also create a *π*-stacking interaction with the ligand, which might aids in stabilizing the ligand enroute to binding pocket. The two residue barriers are very close to binding pocket even F99 and F132 are part of binding pocket itself (Figure 2B). Ligand while adjusting through these residue barriers closely resembles the bound state in terms of distance from the binding pocket, but not in terms of orientation.

To further validate the computational prediction of proposed recognition pathway and especially the role of encounter complex and the residue barriers in sensing mechanism, we rationally designed a set of site-specific mutants and characterized their phenol sensing ability by isothermal titration calorimetry (ITC) studies. The simulation trajectories predicted the first barrier by hydrophobic residues M100, V116 and L119. We surmised that presence of polar residues, instead of these hydrophobic residues, would have formed hydrogen bond across pocket gate residues or with ligand, thereby restricting ligand entry and encounter complex formation. Accordingly, to verify if the phenol-sensing indeed would be controlled by these hydrophobic residues, as posited by simulation, mutations were designed by substituting pocket-gate/channel residues by polar residues (M100Q L119N and M100Q V116N). The mutants of sensor domain (residues 1-229) of MopR were expressed in *E. Coli* and tested for binding to phenol. Mutations did not destabilize the overall structure as indicated by circular dichroism (CD) curves and mutant simulations did indicate hydrogen bond formation (Figure S4). Interestingly, double mutants of MopR in the form of M100Q V116N (*K_d_*=4.35*±*0.07 *µ*M) or M100Q L119N (*K_d_*=2.7*±*0.5 *µ*M) significantly decreased the phenol binding affinity (Figure 3E, S5), in comparison with the wild type MopR (*K_d_*=0.46*±*0.06 *µ*M). On the other hand, a control hydrophobic substitutions (M100V L119V) retained the binding affinity (*K_d_* = 0.45*±*0.06 *µ*M) of wildtype (Figure 3F), indicating that an equivalent hydrophobic gate, similar to M100, V116 and L119 would also facilitate the phenol entry. In order to ascertained the binding events observed through ITC, blank titration experiments were performed wherein the heat of dilution of mutant proteins were negligible (Figure S6) inferring true ligand recognition.

Further, as an interesting proof of concept, the energy cost to open the pocket-gate or channel was enhanced by forming salt bridges between pocket-gate residues (M100K V116D and M100K L119D). Overall structure of mutant construct remain stable as observed by CD curves and mutant simulations did indicate formation of salt bridge (Figure S4). In these mutant constructs, no detectable binding was observed (Figure 3G). Additionally, the second barrier residues (F99, F132) form *π*-stacking interactions with ligand both during binding process and in bound state, if mutated to non-aromatic leucine residues also significantly decrease the binding affinity of MopR (Figure S5). Together, the computational and experimental observations indicate that the ligand approached the buried cavity of MopR via between *α*4 and *β*4 (i.e., pathway-1). In process, the ligand traverse two residue barriers, which are either close to binding pocket (first barrier) or are part of binding pocket (second barrier). Also, the CD curves of mutant constructs and mutant simulations indicate that the structure remain conserved, nevertheless a subtle change in binding pocket cannot be neglected owing to mutated residues proximity to it. All ITC were replicated thrice and for all mutant constructs, the N value maintained at 1 as with wildtype indicating unaltered stochiometry.

Upon examining the sole trajectory that displays the less common binding pathway (pathway-2), it was observed that the ligand’s route through this pathway initially involves ligand entry between *α*4 and *α*3, which causes a significant repositioning of the M105 sidechain. Finally, the ligand crosses between the *α*4 and *α*4 to reach the binding pocket (Figure S7). In any receptor that has an buried binding pocket, an unbound ligand must traverse through a certain width of the protein’s core to access the binding pocket. In the case of MopR, pathway-1 represents the shortest route (measured as the distance between the protein surface and the binding pocket) that the ligand must take to reach the binding pocket (Figure S7). Therefore, we believe that pathway-1 is the most favored pathway for ligand binding in MopR, whereas pathway-2 is either a less significant pathway or an outlier.

### Statistical model reveals four-state ligand-sensing process

A Markov State Model (MSM) was developed to quantitatively characterize the binding process of the ligand-sensing mechanism in MopR. Towards this end, for statistical sampling, a large set of adaptively spawned independent short trajectories were combined with the continuous trajectories, resulting in an aggregate of 25.2 microseconds worth of data. Implied timescale (ITS) analysis of MSM ^30^ predicts that the overall ligand-recognition process by MopR would be optimally described by four key states (Figure 4, S8). The model recapitulates two of these states as ‘phenol-unbound’ (U) and ‘native phenol-bound’ (B) poses of MopR. More importantly, the ITS identifies two short-lived binding-competent metastable intermediates, coined here-after as ‘intermediate-1 (I1)’ and ‘intermediate-2 (I2)’. The ensemble of I1 is represented by phenol waiting outside first barrier (M100, V116, L119), just before the formation of the encounter complex. The recovery of I1 as a non-negligible intermediate indicates that the ligand would spend finite time waiting outside first barrier before this gate opens either spontaneously or after being induced by the phenol. The other key intermediate I2 appears in between the formations of encounter complex and native-bound pose. In this macrostate, phenol is located within the hydrophobic core of MopR and is surrounded by first (M100, V116, L119) and second (binding pocket residues F99, F132, H106) barrier residues. (Figure 4C).

**Figure 4:**
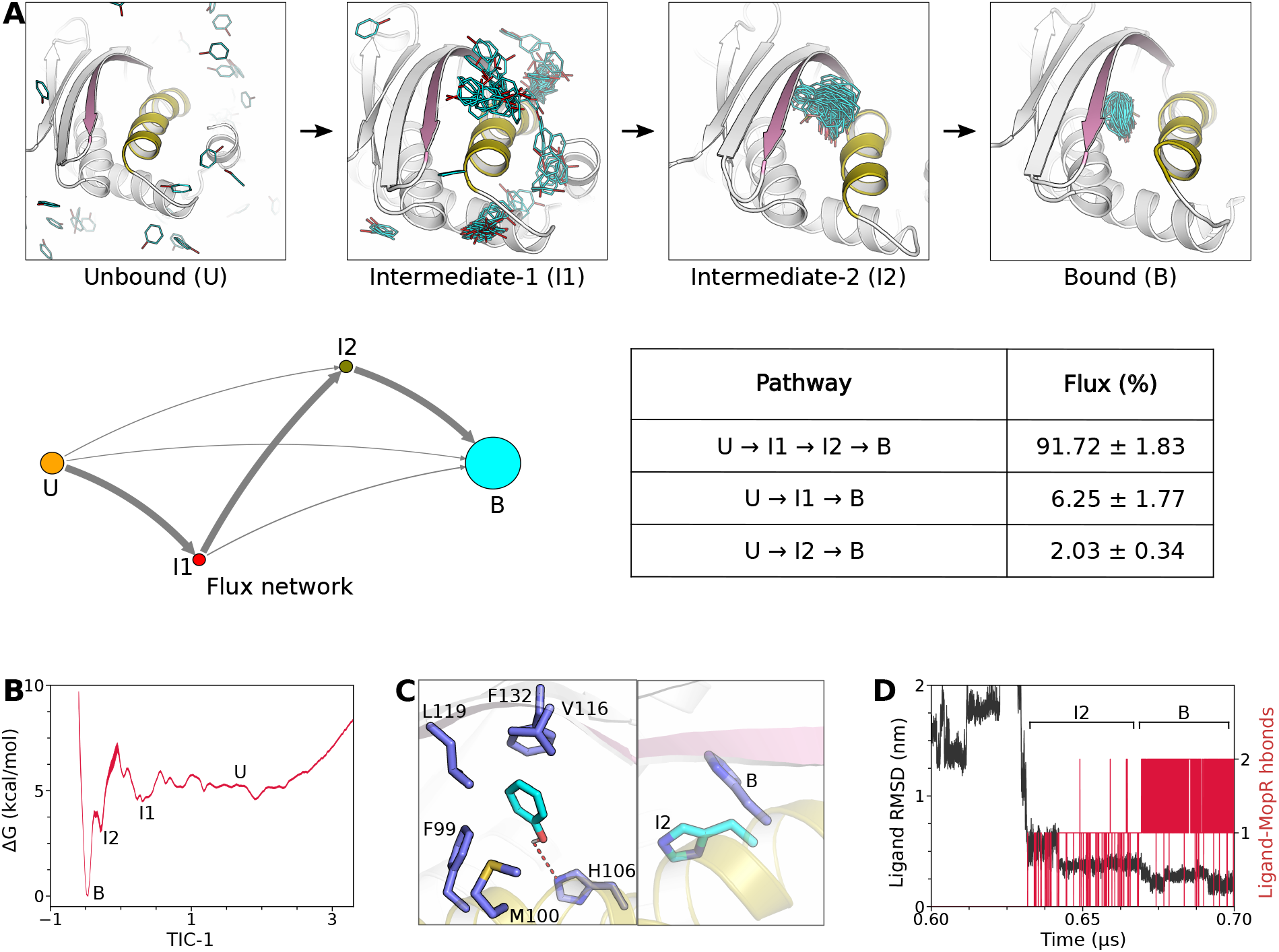
The intermediates. (A) 4 states MSM of binding mechanism in MopR. (B) Free energy plot showing minimas corresponding to 4 metastable states of MSM. Width of the curve represents the error bar. (C) Zoomed residue level view of intermediate-2. (inset) The two states of H106 in I2 and B states. (D) The hydrogen bond time profile of ligand with H106 (I2 and B states) and W134 (only B state).

A quantitative characterization of transition paths between ‘unbound’ and ‘native-bound pose’ ascertained an intermediate-guided pathway i.e., Unbound *→* I1 *→* I2 *→* Bound as the dominant recognition pathway (91.72 *±* 1.83 % probability). The dominant pathway, as predicted by MSM, is akin to aforementioned ‘pathway-1’ involving two barriers. The MSM also predicted a non-zero probability (6.25*±* 1.77 %) for recognition to take place without intermediate-2. This is supported by the fact that out of 12 binding events through pathway-1, 3 of them occurred without any detectable intermediate-2 state.

The key intermediate (I2), present between the first and second barriers with only subtly different from bound state, form *π*-stacking with second barrier residues (F99, F132) and weak hydrogen bond with another binding pocket residue H106 (Figure 4C). The I2 has committor probability of 0.94 *±* 0.02 (Figure S8), indicating that once in I2 state, the ligand has probability of 0.94 to attain successful binding. H106 which form hydrogen bond with ligand in bound state and also in I2 (Figure 2B, 4C) state discern important aspects of ligand sensing in MopR. H106 exist in two different states in I2 and B states which allow it to form hydrogen bond with ligand initially in I2 state and potentially drive the ligand to bound state, where ligand can form hydrogen bond with W134 also (Figure 4C,D, S8). This shifting of H106 marks the beginning of downstream signalling as explained in our previous work.^31^ Additionally, involvement of H106 in I2 state also re-explain our previous observation regarding the pronounced effect of H106 as compared to W134. In particular, H106 (H106A *K_d_*=7.86 *±* 0.01 *µ*M) and W134 (W134A *K_d_*=2.87 *±* 0.01 *µ*M) form hydrogen bond with bound phenol, but have different contribution to binding affinity.^13^ H106 forms hydrogen bond with ligand both in I2 and bound state, while W134 forms hydrogen bond with ligand only in bound state, which explains the pronounced effect of H106 and importance of I2 state. Therefore, we believe this histidine residue is the final selectivity filter 3 which pulls the ligand into the binding pocket and allows it to enter the final bound pose.

### Mechanism of rejection of non-ligands such as substituted bulky phenols and benzene by MopR

In addition to exhibiting high sensitivity toward analyte, a biosensor also needs to be selective. MopR is known to be highly sensitive toward its cognate ligand phenol. However, more importantly the sensor domain of this protein is strategically selective such that it exclusively senses phenol with the highest binding affinity (*K_d_* = 0.46*±*0.06 *µ*M), while larger phenolic derivatives and non-phenolic smaller-sized ligand benzene show significantly decreased affinity (Figure 1, S9).

The phenol-recognition mechanism by MopR, as elucidated in the current investigation, provides a very compelling interpretation of its ligand-selectivity. The aforementioned binding mechanism as well as complementary site-directed mutations had indicated that the pocket-gate created by M100, V116 and L119 residues in *α*4 and *β*4 in MopR can serve as crucial selectivity filter on the way to ligand-entry in the native pocket of MopR. The subtle increase in the M100-V116 distance (either via ligand induction or spontaneous fluctuation) creates a crucial channel (Figure 5) which paves the way for phenol entry. Unfortunately, the residue based M100-V116 distance cannot be correlated with ligands sizes or other metrics. Therefore we estimated the average diameter of phenolic ligands (ligand size) and average diameter of M100-V116 transient channel (channel size) (Figure S9 for estimation method), which can be directly compared to understand size based ligand selectivity. The calculated ligand and channel sizes perfectly correlates with size based binding affinity decrease and M100-V116 distance respectively (Figure 5C, S9). A comparison of various ligand size with that of the transient channel suggests that phenol shall require channel opening of ∼6.67 °*A* or 1.2 nm of M100-V116 distance (Figure 6C). This represents the maximal opening observed in simulations, indicating that the size of the channel would be only sufficient to allow phenol entry (Figure 6C, D). Any bulkier phenol derivative would find it difficult to pass through this size selective channel opening, thereby decreasing the binding affinity. The results indicate that the size based ligand screening occurs at first barrier corresponding to intermediate-1 state.

**Figure 5:**
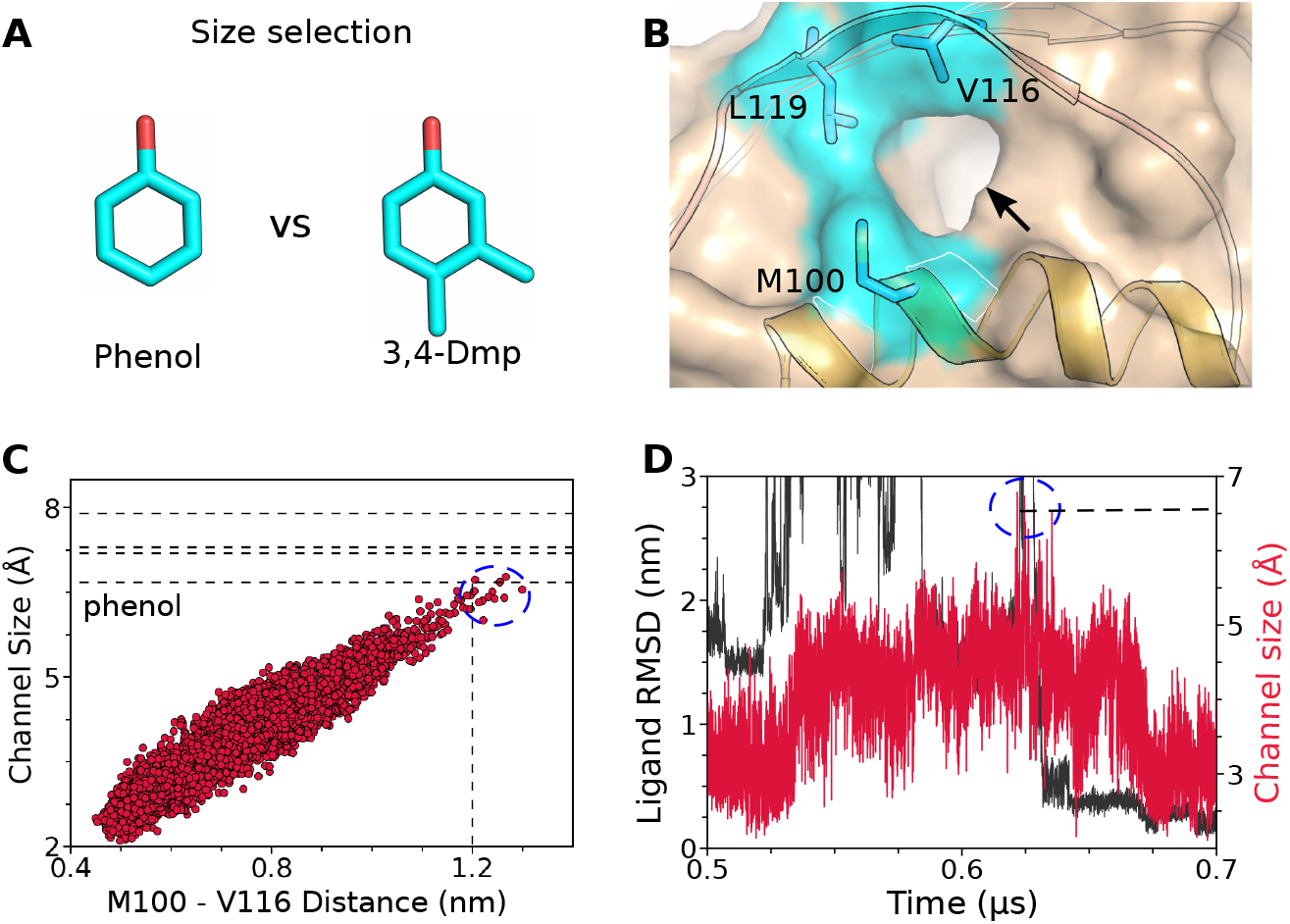
Size based selection. (A) representative phenolic ligands depicting size based ligand screening. (B) The putative size selective channel responsible for filtering the ligands based on size. (C) Channel size and ligand sizes with respect to M100-V116 distance. Dashed line indicates ligand sizes, corresponding to (bottom to top) phenol, o-cresol, m-cresol and 3,4-dmp. (D) Channel size during phenol binding optimal for phenol entry.

**Figure 6:**
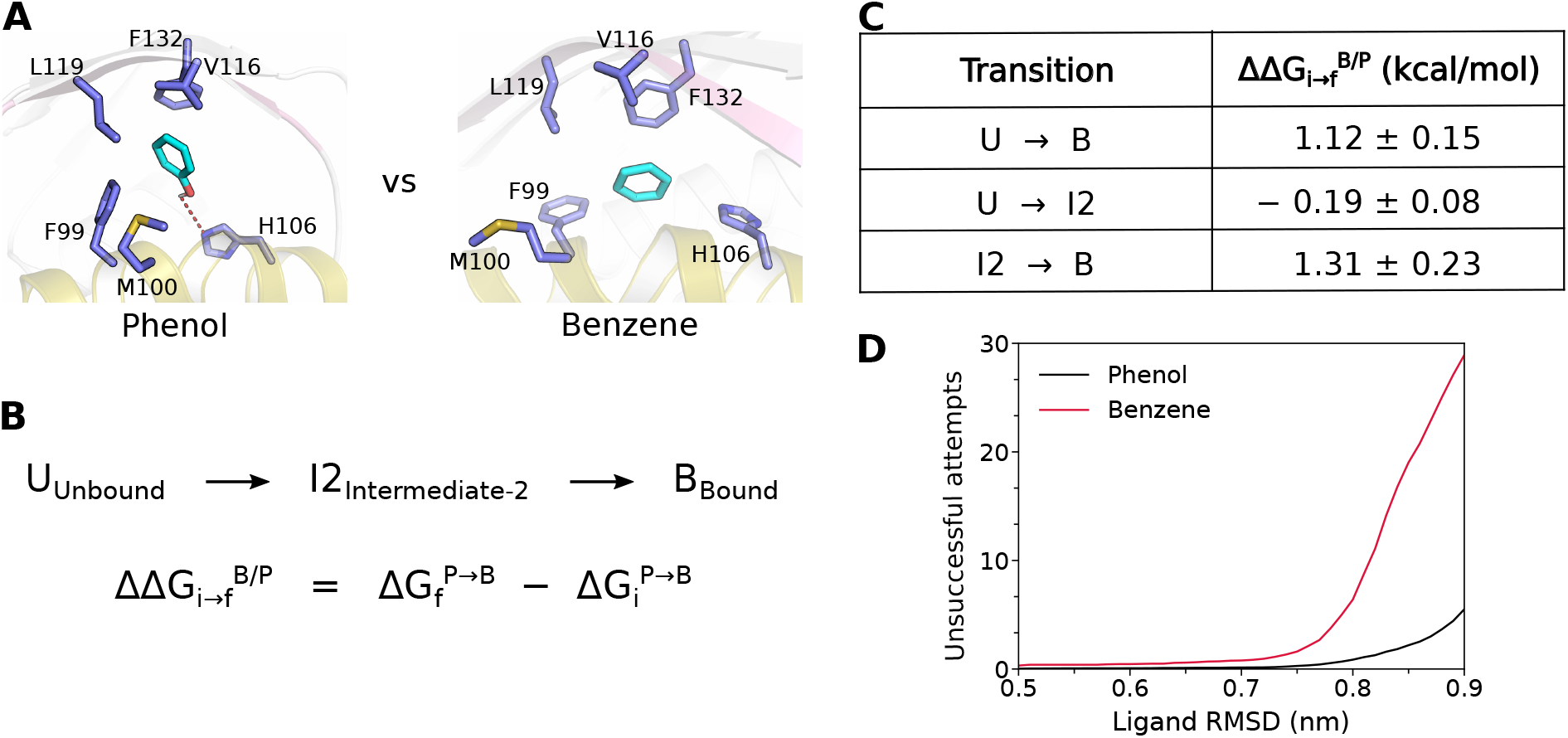
Phenol based selection. (A) Phenol vs benzene in intermeidate-2 state. (B) Relative binding affinity of benzene with respect to phenol for transition i to f, using alchemical thermodynamics (Figure S10 for derivation). (C) FEP/REST relative transition affinities of benzene with respect to phenol for different transitions. (D) The number of unsuccessful binding attempts per microsecond (unbinding after reaching intermediate-2 state [RMSD 0.5 - 0.9 nm]) for phenol and benzene.

The above analysis of size-based ligand selection is under the assumption that larger phenolic ligands would attempt binding to MopR using the same major pathway as phenol, albeit at a significantly decreased success rate than that of phenol. To rule out the possibility that larger phenolic ligands can exercise different sensing mechanism than phenol, similar binding simulations of 3,4-dimethylphenol (3,4-dmp) with MopR were performed. It was found that 3,4-dmp binds at a significantly decreased success rate (one successful event out of three independent simulations) through same pathway as that of phenol (Figure S9), thereby suggesting that the recognition mechanism would be conserved in which ligand-size would impart selectivity.

In addition to being size selective, MopR is also known to be poor sensor of non-phenolic ligands such as benzene even if smaller than phenol in size. Significantly decreased affinity of benzene is interesting as, (i) benzene is smaller than phenol and (ii) orientation dependent ligand entry shall not be required for benzene. Given this logic, this would indicate that ligand entry and formation of encounter complex must be more facile for benzene given that benzene attempts binding through same mechanism. This apparently contradicts the aforementioned argument based on size-selectivity-filter. To understand phenol-based benzene selection, relative transition affinities of benzene with respect to phenol were calculated using FEP/REST^32^ for three transitions i.e., unbound *→−*intermediate-2, unbound *→−* bound and intermediate-2 *→−* bound (Figure 6). We find that in agreement with previous size-selectivity based argument, unbound *→−* intermediate-2 transition is indeed relatively more favourable for benzene (-0.19 *±* 0.08 kcal/mol) than phenol. However, further transition from intermediate-2 bound state was found to be less favourable for benzene i.e., 1.31 *±* 0.23 kcal/mol (Figure 6), combinedly deeming overall benzene binding relatively unfavourable than phenol. In other words, our free energy analysis predicts that benzene shall be able to form encounter complex and attain intermediate-2 state, but would return from intermediate2 state, resulting in more unsuccessful binding attempts, potentially due to lack of hydrogen bonding partner with H106 at intermediate-2 state.

To rule out the possibility that benzene might have alternate sensing mechanism, binding simulations with benzene were performed. Since benzene shows little or negligible binding and to compare the benzene simulations with phenol simulations, benzene binding simulations were performed at significantly high concentration i.e., 0.019 M equal to phenol concentration in binding simulations. It was found that benzene also attempted binding through same pathway-1 (Figure S10). However, comparing benzene and phenol binding simulations reveal that benzene would reach out the intermediate I2 state and but deflect back to solvent (i.e., unsuccessful binding attempt) significantly more often than phenol (Figure 6C, S10). This was in complete agreement with FEP/REST prediction, that phenol-based ligand selection occurred at intermediate-2 state and benzene undergo large number of unsuccessful binding attempts. This indicates H106 controls phenol-based ligand selection, which is in agreement that H106 is also present in other phenolic pollutant sensor DmpR (H100) and HbpR (H108), while not in benzene sensor XylR (Figure 7).^10,14,33^

**Figure 7:**
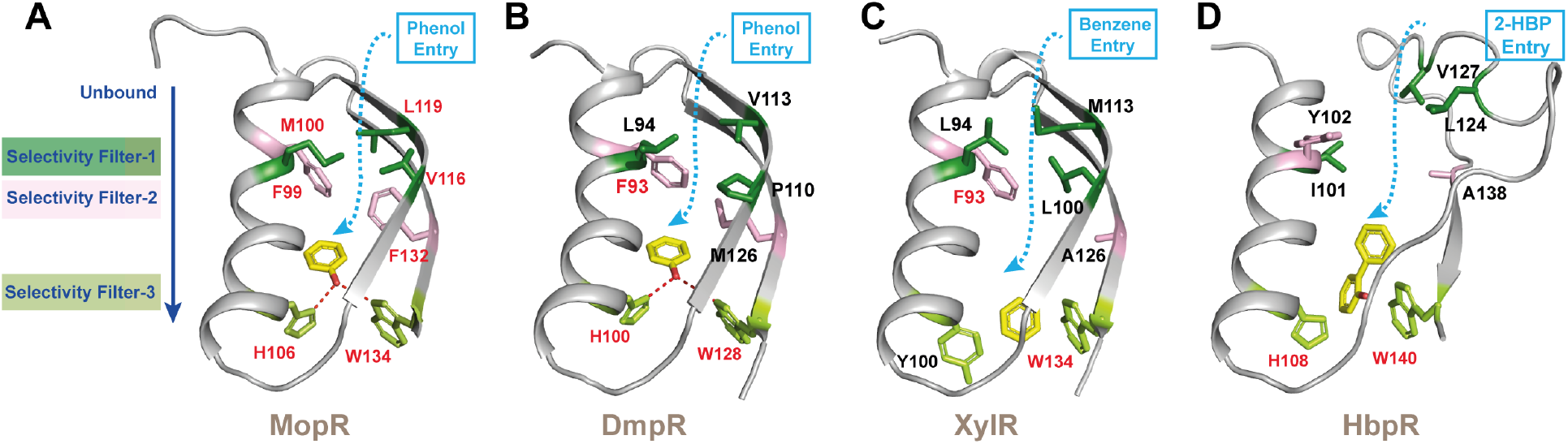
**Comparison of the ligand binding path in aromatic sensing proteins**: (A) A set of three selection filters are conserved in all aromatic sensor units. The pocket sensing residues, selectivity filter 1 are depicted in forest green; selectivity filter 2 in light pink; and selectivity filter 3 H106, W134 in limon green. Schematic representation of ligand transport pathway is highlighted as blue dotted arrow. While coordinates of (A) and (B) for MopR and DmpR are extracted from crystal structures bound with ligand, PDB Id 5KBE and 6IY8 respectively. (C) and (D) depicts, XylR and HbpR which are benzene and hydroxyl biphenyl sensors respectively. Their coordinates are extracted from model generated via Alphafold^36^ and the ligand is docked in the probable binding pocket using Autodock. ^37^ All carbon atoms of ligand are colored in yellow with oxygen atom shown in red.

## Discussion

Bacterial transcription is a tightly regulated process, especially the one controlled under the σ54 RNA polymerases as the genes under this promoters are turned on only under specific external stimuli.^1^ Nature, has therefore devised sophisticated sensing mechanisms to interface with the external environment to avoid mis-regulation or unwarranted cross-talk. In this regard we delve into the mechanism of sensing of MopR, a phenol sensor. Our studies reveal that recognition of the ligand is a complex process, unlike the conventional lock-key hypothesis,^34^ here recognition is not limited to the substrate binding pocket. Rather we believe for buried pockets it starts when the ligand encounters the outer periphery of the protein or binding pocket. In an effort to maintain stringent selection phenol has to encounter three layers of selection to finally reach the buried phenol binding pocket. Here, it is noteworthy to point that during the simulation MopR sensor domain never opens via a dramatic conformational change rather subtle interactions with the pocket-gate help in getting the ligand into the pathway. The first filter, pocket-gate lies near the surface of the protein (between *α*4 and *β*4) and ensures the correct size ligand which is a mono-aromatic group, in the case of MopR, enters the protein matrix (selectivity filter 1). This filter is exposed to a milieu of molecules and therefore a critical selection juncture. Analysis of other proteins that harbor similar sensor domains such as DmpR, XylR, HbpR etc. show that a pocket-gate filter is a common feature and has mostly aliphatic hydrophobic amino acids such as methionine, leucine valine etc. ^10,35^ (Figure 7) XylR,^33^ a benzene sensor mostly closely resembles the pocket gate organization of MopR and this is one of the reasons in the simulation’s benzene is able to pass the first selection point. Comparison of this region of MopR with DmpR shows that the pocket evolution is subtly tuned into the protein sequence. In DmpR, a proline has been replaced in *β*4 instead of a valine, which then introduces a kink in this region, facilitating the bulkier ligand 2,3-dimethyl phenol to enter. Interestingly, in HbpR which is a hydroxy bi-phenyl sensor the pocket filter has a similar hydrophobic configuration but a wider pocket to accommodate the biphenyl-ring recognition. Thus, a fine interplay and placement of apt residues at the selectivity filter 1 stage controls the size and nature of the ligand that enters a particular sensor module.

The second filter in the MopR phenol sensor helps direct the ligand towards the binding pocket and primes it for selection to the next filter. A comparison of the four available sensor modules depicts that helix *α*4 almost always harbors a phenylalanine with the corresponding amino acid at the *β*4 end adjusting according to the size of the entering ligand. ITC studies in conjunction with the comparison as depicted in Figure 7 support this hypothesis and reasserts that it is *β*4 residue (F132 in MopR) that evolves as per the ligand requirement. However, the most crucial filter is the rejection of isostructural hydrocarbons such as between phenol and benzene for the phenol sensor which is orchestrated via a key histidine residue (final selection filter 3). Simulations show that in MopR, absence of interaction of benzene with H106 destabilizes the encounter complex leading to it’s rejection. All aromatic phenol group sensing proteins harbor the histidine residue however, for proteins that accept only hydrocarbon moiety such as XylR (benzene sensor) this histidine is missing (Figure 7). Moreover, previous reports through protein engineering studies show that where this histidine is replaced by alanine, phenol sensing is completely abolished.^13^ Whereas, in MopR it has been shown that replacement of histidine to tyrosine H106Y helps creates an effective benzene sensor and leads to complete rejection of phenol.^17^ This highlights the importance of this filter that selects for phenols. Moreover, HbpR which identifies hydroxy bi-phenyls also has this histidine sensor residue conserved, thereby reasserting it’s central role in aromatic sensing (Figure 7).

Enzymes with buried pockets have various ways to allow ligands inside, either via opening/closing of a specific tunnel, or via an allosteric switch that opens up a transient path or via formation of temporary pockets that then facilitate opening of the main binding site or via breathing motion that allow side chain fluctuations making way for the ligand. ^22,26,27,38^ The MopR simulations have revealed that the system allows the sensor molecule to enter in a seemingly passive fashion with fluctuations in the protein being the primary way in which the ligand approaches the protein. Once the entry process is enroute it then then subjects the ligand to a stringent selection. One would argue the need of this stringency in selection. In fact, it appears that concentration of the small molecule/pollutant acts as a descriptor of the external stress. The processes initiated by recognition of the small sensor molecule by the sensor module are quite energy intensive to the cell. Therefore, a judicious decision-making strategy at this early juncture is paramount. For instance, in MopR, the sensor unit which is linked to the ATPase unit gets turned-on upon binding of phenol and subsequently triggers σ54 RNA polymerase that then transcribes a substantial set of phenol degrading genes and shifts the whole energy requirement of the cell from glucose to phenol, an energy intensive process. Since no chemical reaction is taking place, it is the sensor module that needs to maintain the stringency as there are no further barriers once the ligand reaches the sensing pocket. Therefore, the whole sensing process has been layered under selection filters to avoid accidental triggering of the pathway.

In conclusion, here the described understanding of the binding pathway helps in not only understanding the basic mechanism behind ligand selection but opens new avenues for intelligent biosensor design for this important class of aromatic pollutant sensors.^39–42^ Since the multi-step selection process takes place before the ligand actually reaches the binding pocket, an important aspect to consider while designing biosensors rests on tweaking parameters that govern the propensity of entry of a particular ligand.^43,44^ New ligand-biosensor combinations can be created by altering the residues that line the entry path such that only the molecule of choice is allowed to reach the binding pocket. The multi-layered selection process can be extended to benchmark the determinants necessary to achieve optimal protein-ligand design.

## Methods

### Computational methods and model

#### Unbiased binding simulations

The crystallographic structure of wildtype MopR dimer in phenol bound form (PDB code: 5KBE) was used as starting point for all our simulations.^13^ The missing residues were modelled using charmm-gui.^45^ The protonation states of titrable amino acids corresponded to neutral pH except for C155, C181 and C189, which were modelled as negatively charged due to coordination to *Zn*^2+^ ion. Each protomer of MopR contained one zinc ion tetrahedrally coordinated to C155, E178, C181 and C189. Cysteine sidechain pKa was assumed to shift in presence of close zinc leading to deprotonation^46^ and trial simulations showed stable zinc conformation with deprotonated cysteines (Figure S11). For binding simulations, the phenol-free apo form of the MopR dimer was placed at the center of octahedron box with 10 °*A* minimum distance between protein surface and box and empty space filled with water and ions. The system was solvated with around 18500 water molecules and sodium and chloride ions were added to keep the sodium chloride concentration at 200 mM and overall neutral charge. Ten phenol molecules, corresponding to experimentally employed 1:5 protein-ligand ratio^13^ were placed at random positions in the simulation box such that no phenol molecule was in contact with MopR dimer. The system thus prepared led to phenol concentration of 19 mM which is much less than phenol water solubility limit of 0.88 M. Three such systems with randomly placed phenol molecules were generated and two all atom MD simulations were started from each of the systems. The total system overall included around 63000 atoms. The protein, phenol and ions were parameterized with Charmm36m forcefield^47^ and charmm-TIP3P water model^48^ was used. All MD simulations were performed with Gromacs 20xx packages benefitting with usage of graphics processing Units. The simulations were performed at average temperature of 298 K as per experimentally employed condition, and 1 bar pressure. The temperature was maintained using Nose-Hoover thermostat^49,50^ with a relaxation time of 1.0 ps and pressure was maintained using Parrinello-Rahman barostat^51^ with a coupling constant of 5.0 ps. The long range electrostatic interactions within 1.2 nm cut-off were calculated with the use of Particle Mesh Ewald summation^52^ and Lennard Jones interactions within 1.2 nm cutoff were calculated using Verlet cutoff scheme.^53^ All the hydrogen bonds were constrained using LINCS algorithm^54^ and water hydrogen bonds were fixed using SETTLE algorithm.^55^ All equilibration and production runs were performed at 1 and 2 fs timesteps respectively. 12 replicates of unbiased all atom MD simulation trajectories were spawned, each preceeded via 500 ps restrained NVT equilibration and differing in initial protein-phenol configuration and initial velocities as per protocol of our previous works. ^26,27^ The binding simulations ranged between 1 to 4 microseconds with a total simulation length of 20 microseconds. The simulations were stopped only after any of the phenol molecule(s) had remained bound for sufficiently long duration (*>* 200 ns) to any one or both the protomer(s) of MopR dimer. The binding process at any instant was checked by two metrics; (i) center of geometry distance between phenol and binding pocket or inner shell as defined in this work and (ii) the RMSD of any phenol molecule at an instant in simulated conformation relative to the crystal structure (PDB id: 5KBE) of phenol-bound MopR. The RMSD cut-off value of 3 °*A* was used to confirm the binding event. Apart from the binding simulations, apo and bound forms of MopR dimer were individually sampled (apo and bound simulations) using 1 microsecond long simulations, following the same procedure as above. Further, binding simulations with 3,4-dimethylphenol and benzene were performed, following the same procedure as for phenol. 3,4-dimethylphenol and benzene were parametrized using charmm36 forcefield parameters. Additionally, simulations of mutant MopR were also performed for M100K L119D, M100K V116D, M100Q L119N and M100Q V116N mutations. The in-silico mutations were performed using charmm-gui.^45^ For each mutant, 3 independent simulations each of 200 ns timelength were performed.

#### MSM analysis

For binding pathway analysis, a Markov state model (MSM)^56,57^ was built using PyEMMA python library. ^58^ For building a comprehensive MSM, in addition to aforementioned unguided aggregated binding trajectories (both pathway-1 and pathway-1), extenstive adaptive sampling was performed yileding a total of 25.2 *µ*s worth of data. For adaptive sampling, to improve the simulation statistics, a large set of (122 total) short independent MD simulations (each 100 ns long) were initiated from various binding competent states as characterised in long binding simulations. For this purpose, starting structures were extracted from binding simulations representing bound to unbound spectrum of ligand states (ligand RMSD = 0.1 - 1.8 nm). Both the aforementioned metrics (protein-ligand distance and RMSD) which were used to detect phenol-binding in long unguided trajectories, were also employed to construct the MSM. The 2 dimensional input data was linearly transformed into slow linear subspace by time-lagged independent component analysis (TICA)^59^ to 1 TIC component using a correlation lag time of 5 ns, covering more than 95% of kinetic variance. The one-dimensional TICA output was further discretized to 300 microstates using k-means clustering algorithm.^60^ The resulting micro states were used to construct the final MSM at a lag time of 250 ps. The MSM were further coarse grained to 4 metastable macrostates based on ITC plots, using PCCA algorithm.^61^ Finally, the transition path theory was applied to analyze the transition pathways and path fluxes among the macro states. ^62^ The MSM protocol was repeated for 120 replicates with one to three random trajectory datasets were removed for each iteration. The errors were calculated as standard deviation in 120 iterations.

#### Free energies of pocket gate opening

Umbrella sampling approach^63^ was employed to map free energy for M100-L119 pocket gate opening that was found to be crucial for ligand binding. The visual inspection of the MD trajectory had indicated that the pocket gate opening was mostly triggered by side chain movement. Accordingly, the center of mass (com) distance between side chains of M100 and L199 was used as collective variable (CV) for projecting the free energy profile using umbrella sampling. The value of the CV ranged between 0.6 and 1.1 nm representing ‘pocket-gate closed’ to ‘pocket-gate open’ and was discretised into 11 windows with an uniform interval of 0.05 nm. The starting configurations for each umbrellas were picked from unbiased MD trajectories. We derived free energy profile for three pocket-gate opening in three independent scenarios: a) For ‘apo conformation’, pocket-gate opening was measured in absence of phenol and frames were picked from apo simulations; b) for ‘encounter-complex’, pocket gate opening was measured with phenol entering the binding pocket and frames were picked from binding simulations; c) for ‘with bound ligand’, pocket-gate opening was measured with phenol in the binding pocket and frames were picked from bound simulations. Each umbrella windows was equilibrated for 500 ps and sampled for 20 ns ensuring sufficient sampling of entire CV space by restraining each umbrella to desired value of CV using suitable harmonic potential. Finally, the weighted histogram analysis method^64^ was used to reweight the umbrellas to get potential of mean force as a function of CV.

#### Estimation of Relative binding affinities

The binding affinities of benzene relative to phenol were computed using Free energy perturbation(FEP) approach^32^ in combination with Replica Exchange with solute Tempering(REST)^65^ using a similar approach as in our previous works. ^66^ In particular, the relative binding affinities were computed for binding either from unbound to intermediate-2, unbound to bound or intermediate-2 to bound state of the ligand. For this purpose, three different system states were simulated, with hybrid ligand undergoing phenol to benzene transition (i) in bulk solvent representing the unbound state; (ii) in pathway 1 intermediate-2 pose representing the intermediate-2 state, and (iii) in binding pocket representing the bound state. The relative binding affinities of benzene with respect to phenol for transition from state ‘i’ to ‘f’ was calculated using alchemical thermodynamic cycle ^67^ as:

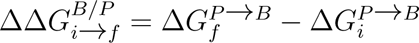

where: 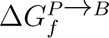 represents free energy change of phenol to benzene transition in state ‘f’ and 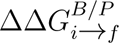 represents the relative binding affinity of B with respect to P while transitioning from state ‘i’ to ‘f’.

In FEP simulations, the initial configurations for intermediate-2 and bound state were obtained from the unbiased binding simulations. The phenol to benzene transition was carried out in 24 discrete steps of coupling parameters *λ* ranging from *λ*=0 (where hybrid ligand represents the phenol) to *λ*=1 (where hybrid ligand represents the benzene). The hybrid ligand configuration and topology parameters were created using pmx scripts,^68^ giving hybrid ligand with 3 possible dummy atoms and much lower similarity score 0.2143, owing to close resemblance between phenol and benzene. To employ the REST with FEP, a hot region was defined for each state of system, (i) For unbound state, phenol was used as hot region, (ii) for intermediate-2 state, along with ligand H106, F99 and F132 represents the hot region and (iii) for bound state, along with ligand H106, W134 and F99 represents hot region. The only residues involved in important protein-ligand interactions in intermediate-2 and bound state were considered for hot region, because large hot region increases uncertainity in energy calculations. A total of 24 *λ* windows were simulated for FEP/REST. For FEP/REST, the temperature of the hot region was raised using appropriately scaled Hamiltonian based of specific *λ* value, with the effective temperature profiles at 298, 337.7, 383.4, 434, 492.7, 559.2, 634.7, 720, 816.5, 926.8, 1051, 1192, 1192, 1051, 926.8, 816.5, 720, 634.7, 559.2, 492.7, 434, 383.4, 337.7, 298 K, ensuring same physiological (298 K) temperature for *λ*=0 and *λ*=1. Each *λ* window, was equilibrated for 3 ns and sampled for 6 ns. For FEP/REST, replica exchange was attempted after every 2000 steps. The final binding free energies were calculated by Bennett Acceptance Ratio method.^69^

### Experimental method and materials

#### Molecular cloning and protein expression

In order to validate the computational results, various mutations were incorporated into the *MopR^AB^* (residue 1 to 229) construct of full length MopR protein of *Acinetobacter calcoaceticus* NCIB8250 that was cloned into a modified pET28a expression vector as in our previous work.^13^ All the point mutations were performed by employing standard site-directed mutagenesis protocol using the Phusion DNA polymerase from New England Biolabs. The reaction mixture compositions were as follows: 1X phusion buffer, 0.2 mM dNTPs, 0.5 *µ*M of forward and reverse primer, 10 ng template plasmid and 0.3 U Phusion DNA polymerase. The programme optimized for the site directed mutagenesis was as follows: Initial denaturation at 95 *^o^*C for 2 min followed by 18 cycles of denaturation (98 *^o^*C for 30 second), annealing ((Tm -5) *^o^*C for 30 seconds), extension (72 *^o^*C for 5 minutes) and then a final extension 72 *^o^*C for 5 minutes. The product was confirmed on 0.8 % agarose gel was digested with Dpn1 for 2 hr at 37 *^o^*C. The Dpn1 digested product was then transformed into E. coli DH5*α* cells and the obtained single colony were processed for plasmid isolation. All the mutations in the obtained clones were confirmed by DNA sequencing. These cloned constructs were transformed into Escherichia coli Rosetta (DE3) cells and grown at 37*^o^*C till OD600 reached 0.6-0.8 followed by induction with 1mM IPTG (isopropyl-*β*-D-thiogalactopyranoside) at 16*^o^*C for 16 hours. The bacterial cells grown were harvested by centrifugation at 4000 rpm for 20 min. All the mutants were expressed as N-terminal His tag fusion proteins.

#### Protein purification

The harvested cells were resuspended in lysis buffer (50mM HEPES buffer, pH 7.5; 2mM Imidazole; 200mM NaCl), lysed by sonication, and centrifuged to separate the debris from the cell extract. The separated cell extract was then mixed with Ni-NTA resin that was pre-equilibrated with the lysis buffer and was subjected to gentle stirring for 1.5hrs. The resin was transferred to a column and was subjected to washing with wash buffer (50mM HEPES buffer, pH 7.5; 30mM Imidazole; 200mM NaCl), and subsequently protein was eluted with elution buffer containing 50mM HEPES buffer, pH 7.5; 200mM NaCl. The eluted fractions were further concentrated and exchanged to buffer containing 25mM HEPES, pH 7.5, 80mM NaCl, 0.5mM DTT using an Econo-Pac 10DG (Bio-Rad, CA, USA) column. The protein fractions obtained were then pooled together, concentrated, flash-frozen in liquid N2, and stored at -80 *^o^*C until they were used.

#### Phenol binding experiment using ITC

The ITC experiments were performed using MicroCal iTC200 (GE Healthcare) to calculate the binding affinity of phenol towards various *MopR^AB^* mutants.^70^ All the protein and ligand (phenol) samples were prepared in buffer (A) containing 25mM HEPES (pH-7.5) and 80mM NaCl. The sample cell containing 10-40*µ*M of *MopR^AB^* mutants was titrated against 100-400*µ*M of phenol. The concentrations of protein and ligand used in different ITC experiments varied as per requirement, in order to attain optimal saturation for a particular titration curve. The volume of the titrant (phenol) added at each injection into the sample cell was 2*µ*l for 5 sec. A range of 16-20 injections was performed for each experiment with an interval of 120 sec between each successive injection. The temperature was maintained at 25 *^o^*C. The stirring rate was kept constant at 750 rpm throughout the ITC experiments. In all the ITC experiments, the ligand was titrated against buffer A and subtracted from the raw data prior to model fitting, in order to nullify the heat of dilution. The data obtained were fitted and analyzed using one set of sites model with Origin 7 software. All the titration experiments were replicated thrice in order to validate the results. The protein concentrations for the various *MopR^AB^* mutants used in ITC were quantified using an UV-VIS spectrophotometer by measuring their absorbance at 280 nm.

