## Supplemental figures for "Identifying Selectivity Filters in Protein Biosensor for Ligand Screening"

**Movie1**: Motion picture showing ligand (phenol, cyan vdw) binding to MopR sensor domain (cartoon) through major pathway-1, i.e., ligand entry in between $\alpha 4$ (tan) and $\beta 4$ (pink). MSM derived metastable). MSM derived metastable states are written on bottom. Pock). MSM derived metastableet - red surface , pock). MSM derived metastableet-gate/channel residues (M100, V116, L119) - stick). MSM derived metastable representation.

**Movie2**: Motion picture showing ligand (phenol, cyan vdw) binding through minor pathway-2, i.e., ligand entry in between $\alpha 4$ (tan) and $\alpha 3$ (purple), $\alpha 6$ (green). Unbound, bound and intermediate states are written on bottom. Pock). MSM derived metastableet - red surface, important residues (D82, T86, M105) - stick). MSM derived metastable representation.

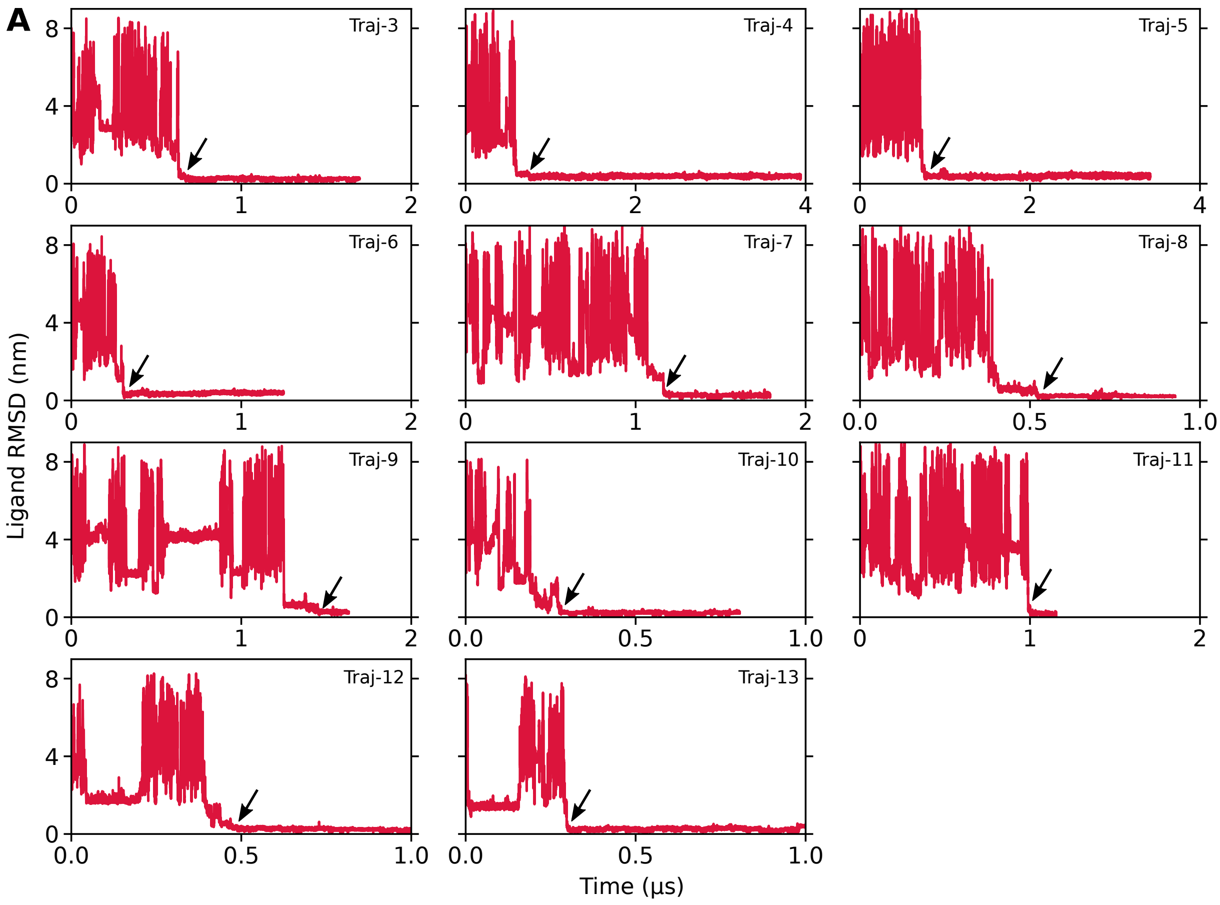


**B**

| Trajectory | Binding Time (μs) | Pathway |
|---|---|---|
| Traj-1 | 0.559 | p1 |
| Traj-2 | 3.221 | p2 |
| Traj-3 | 0.665 | p1 |
| Traj-4 | 0.748 | p1 |
| Traj-5 | 0.771 | p1 |
| Traj-6 | 0.310 | p1 |
| Traj-7 | 1.170 | p1 |
| Traj-8 | 0.520 | p1 |
| Traj-9 | 1.452 | p1 |
| Traj-10 | 0.281 | p1 |
| Traj-11 | 1.001 | p1 |
| Traj-12 | 0.747 | p1 |
| Traj-13 | 0.300 | p1 |

| Binding Statistics | |
|---|---|
| Total bindings | 13 |
| pathway-1 | 12 |
| pathway-2 | 1 |
| $k_{on}$ | $3.5 \times 10^7$ $M^{-1}s^{-1}$ |

**Figure S1:** (A) The RMSD time profiles of binding trajectories Traj3-13. (B) Observed binding time and estimated on-rate constant $k_{on}$ .

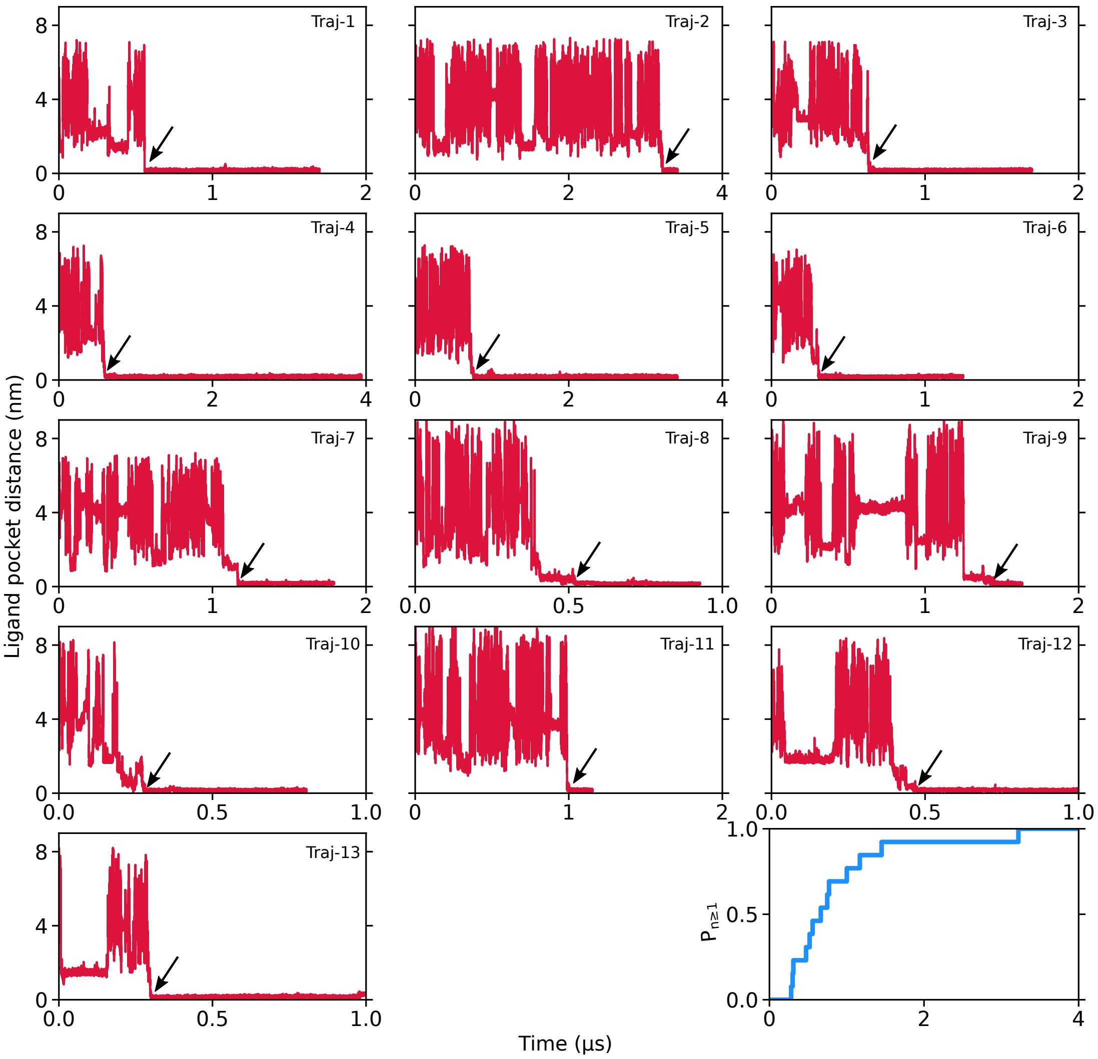


**Figure S2:** Time profile of distance between center of geometry of binding pocket residues and ligand for all binding events (Traj1-13). The empirical probability (blue) of phenol binding to MopR based on observed binding times from simulations. This curve represents the probability of observing one or more binding event within a particular time (given that multiple simulations are performed).

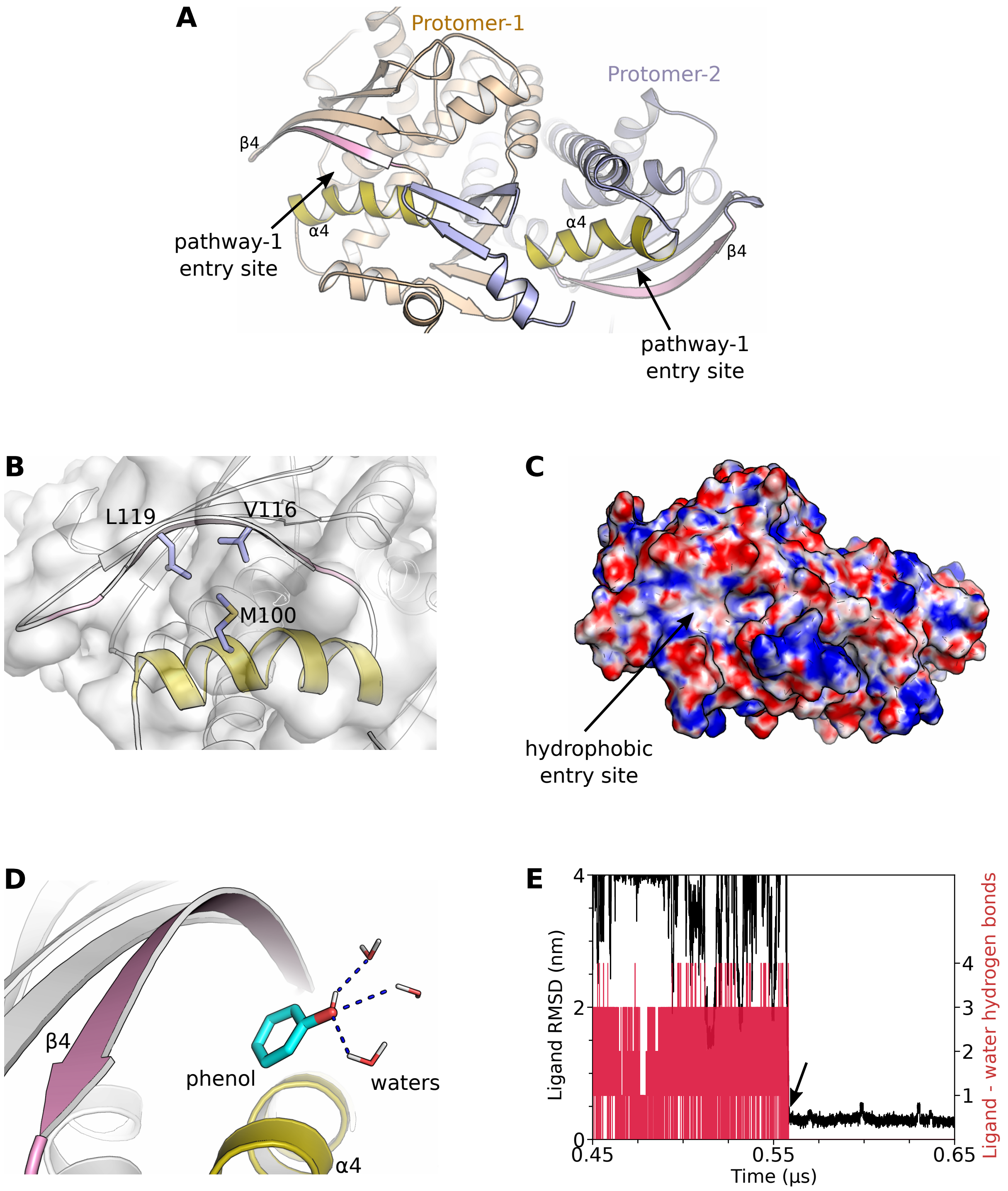


**Figure S3:** (A) Pathway-1 entry site for each protomer lies in opposite direction in dimeric MopR rendering independent ligand binding in each protomer. (B) The first barrier residues in absence of ligand. M100 remains in contact with V116 and L119. (C) Poisson-Boltzman electrostatic surface of MopR. Red to blue spectrum represents postitive to negative potential (± 2 kcal/mol). Pathway-1 entry site represents the hydrophobic surface. (D,E) Orientation specific ligand entry owing to hydrogen bonding between ligand and water molecules. The water-ligand hydrogen bonds break during ligand entry (arrow).

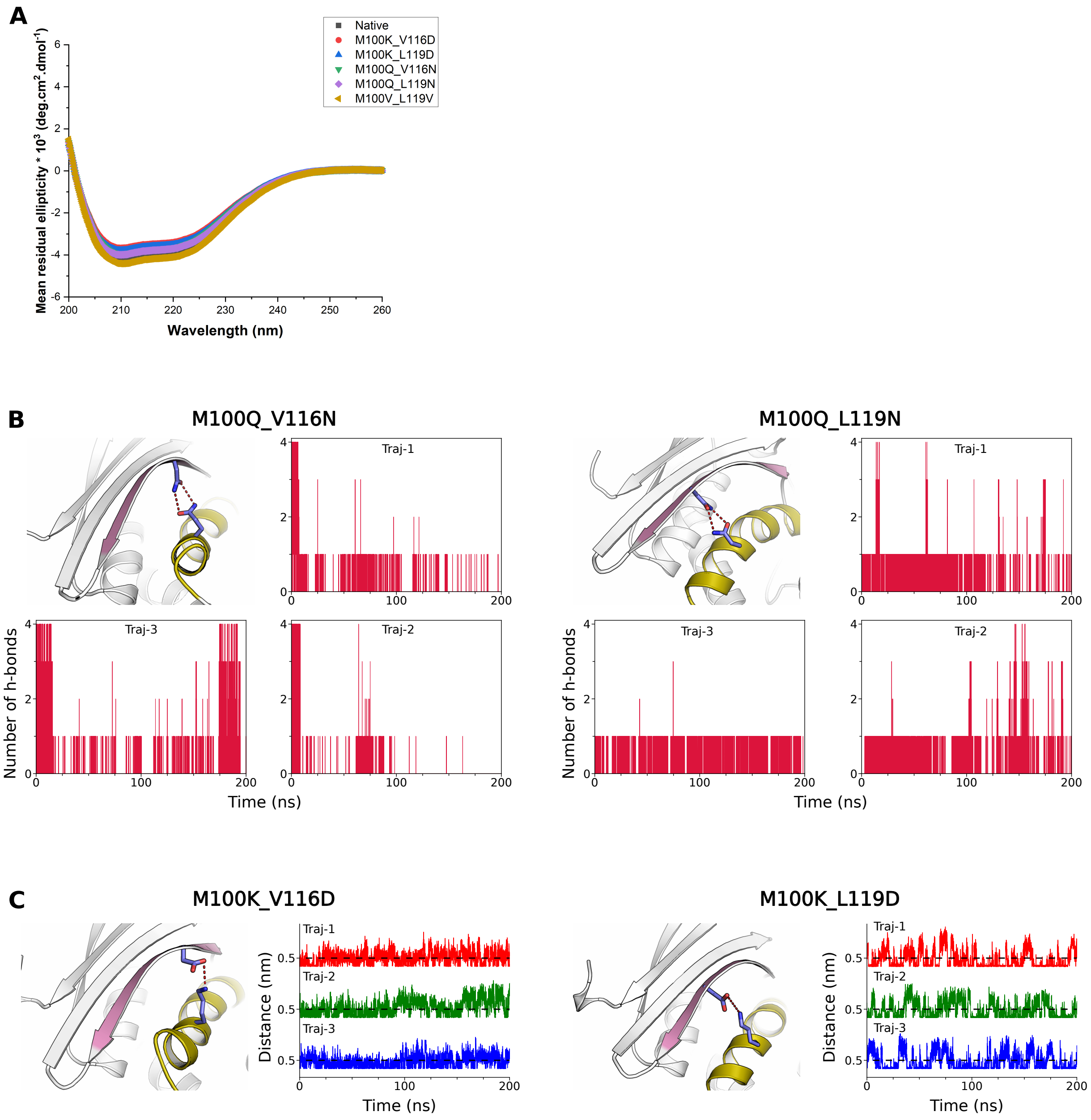


**Figure S4:** (A) Circular dichroism spectra of wildtype and mutant constructs of MopR sensor domain. (B) Hydrogen bond formation in M100Q_V116N and M100Q_L119N mutant simulations. (C) Salt bridge formation in M100K_V116D and M100K_L119D mutant simulations.

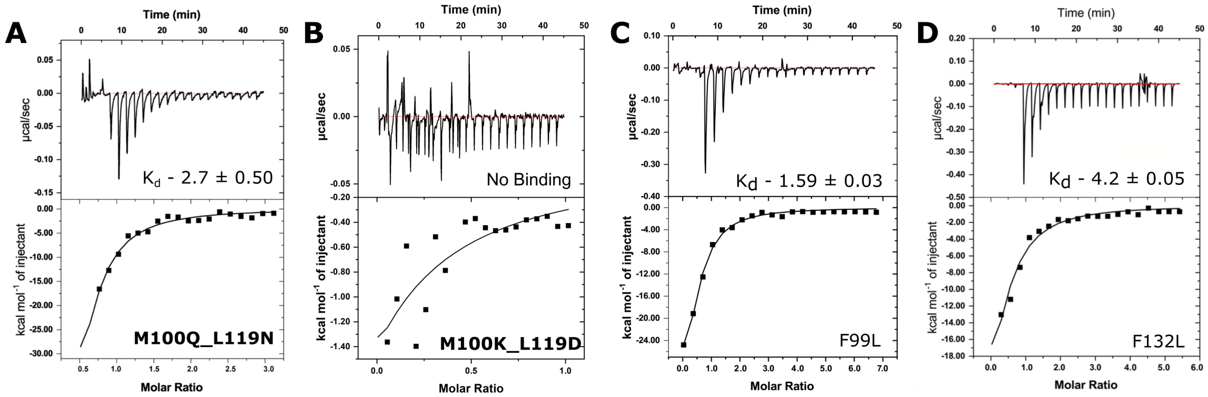


**Figure S5:** ITC binding studies of different mutant versions of MopR. The effect of mutation to a (A) charged polar reside (B) formation of a salt-bridge and to (C-D) non-aromatic leucine residue is shown.

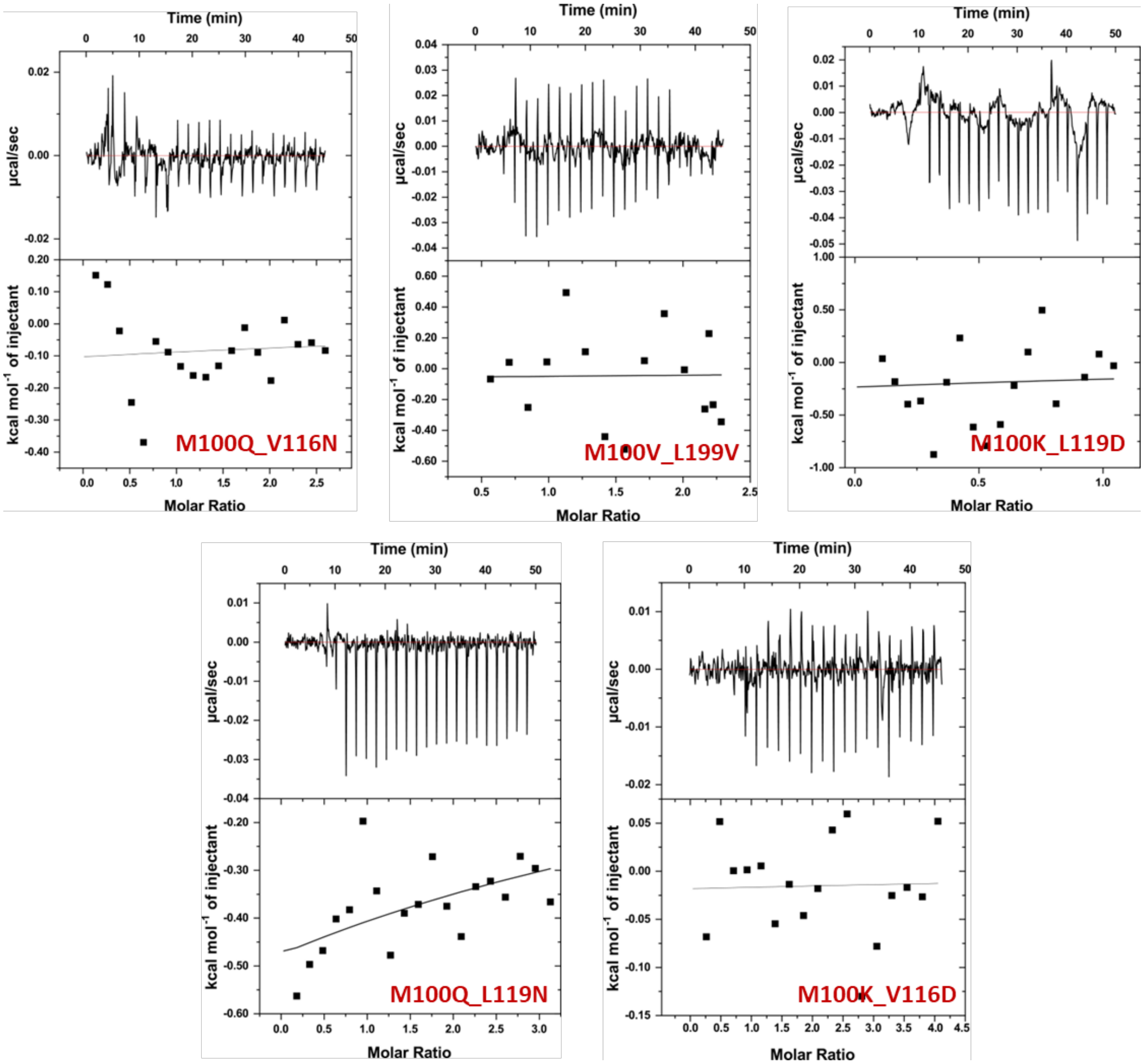


**Figure S6:** Isothermal titration calorimetry experiments showing the heat of dilution for each mutant version of the MopR.

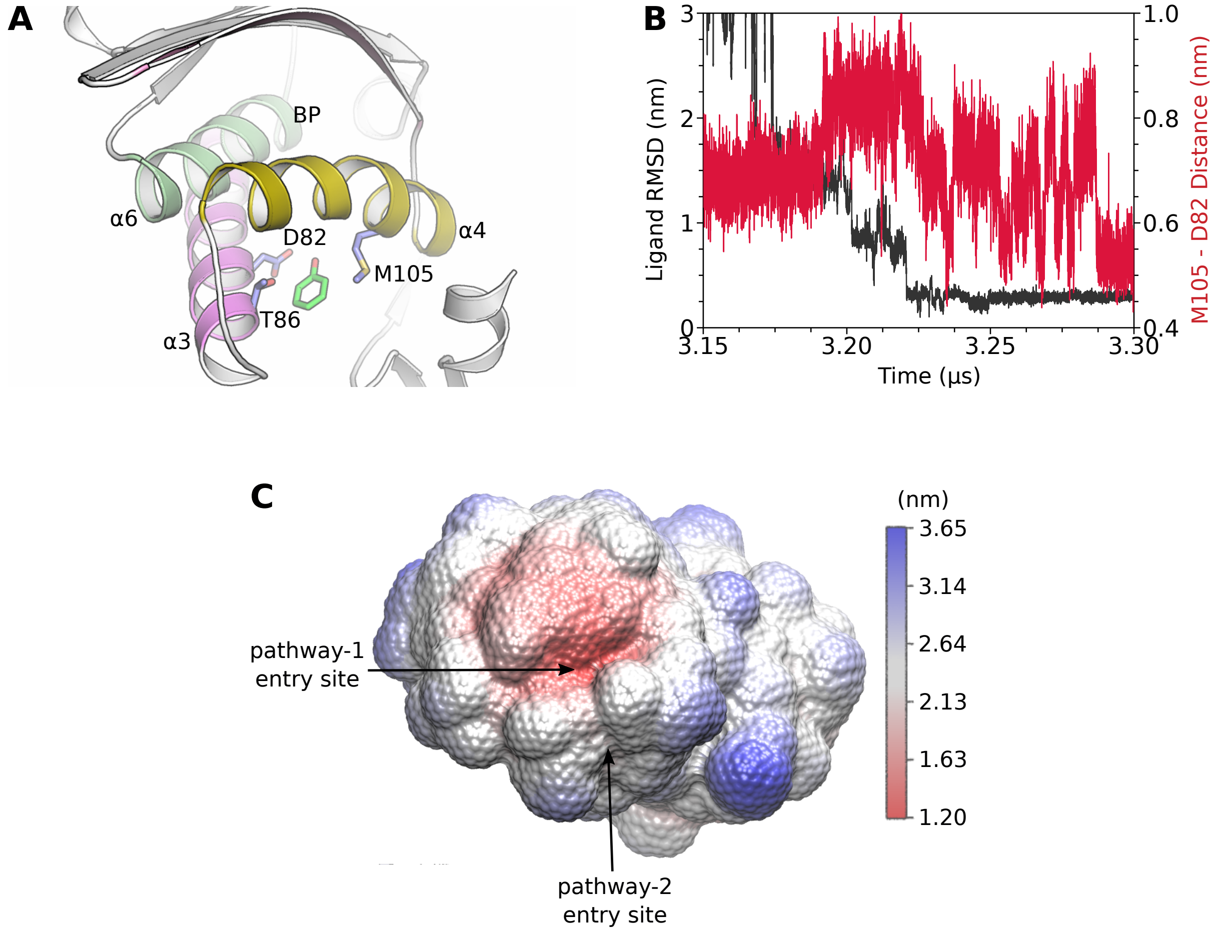


**Figure S7:** (A, B) Ligand entry through pathway-2 (via $\alpha4$ and $\alpha3$) associated with significant movements of M105 sidechain. (C) Distance from the surface (phenol excluded) to the binding pocket. pathway-1 entry site represent the closest entry site or shortest path to binding pocket.

A

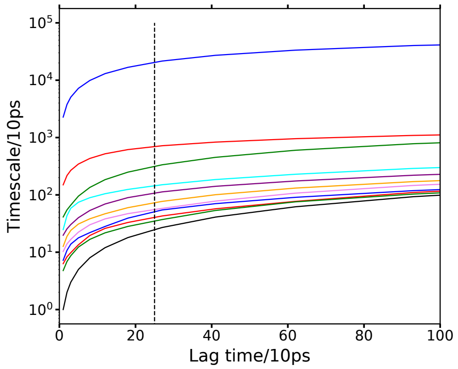


B

| State | Population (%) | Committor probability |
|---|---|---|
| Unbound | 0.00 ± 0.00 | 13.85 ± 0.18 |
| Intermediate-1 | 3.88 ± 0.26 | 0.15 ± 0.02 |
| Intermediate-2 | 4.21 ± 0.54 | 0.94 ± 0.02 |
| Bound | 78.06 ± 0.55 | 1.00 ± 0.00 |

C

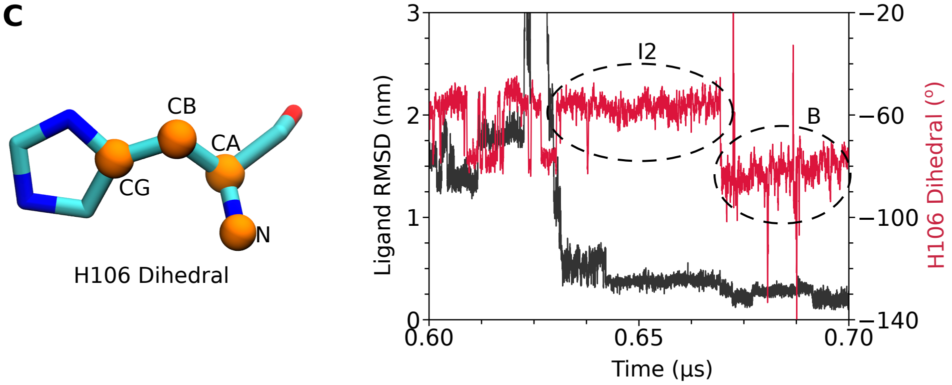


**Figure S8:** (A) (A) The Implied timescale (ITS) plot showing existence of 4 metastable states (blue, red, green). One metastable state is at infinity. Closely separated states at $\lambda = 2$ (red) and $\lambda = 3$ (green) indicates the presence of two intermediate states. (B) Stationary populations and committor probabilities of metastable states. (C,D) Switch in H106 conformation during intermediate-2 to bound state transition as measured by H106 dihedral ($C_G - C_B - C_A - N$).

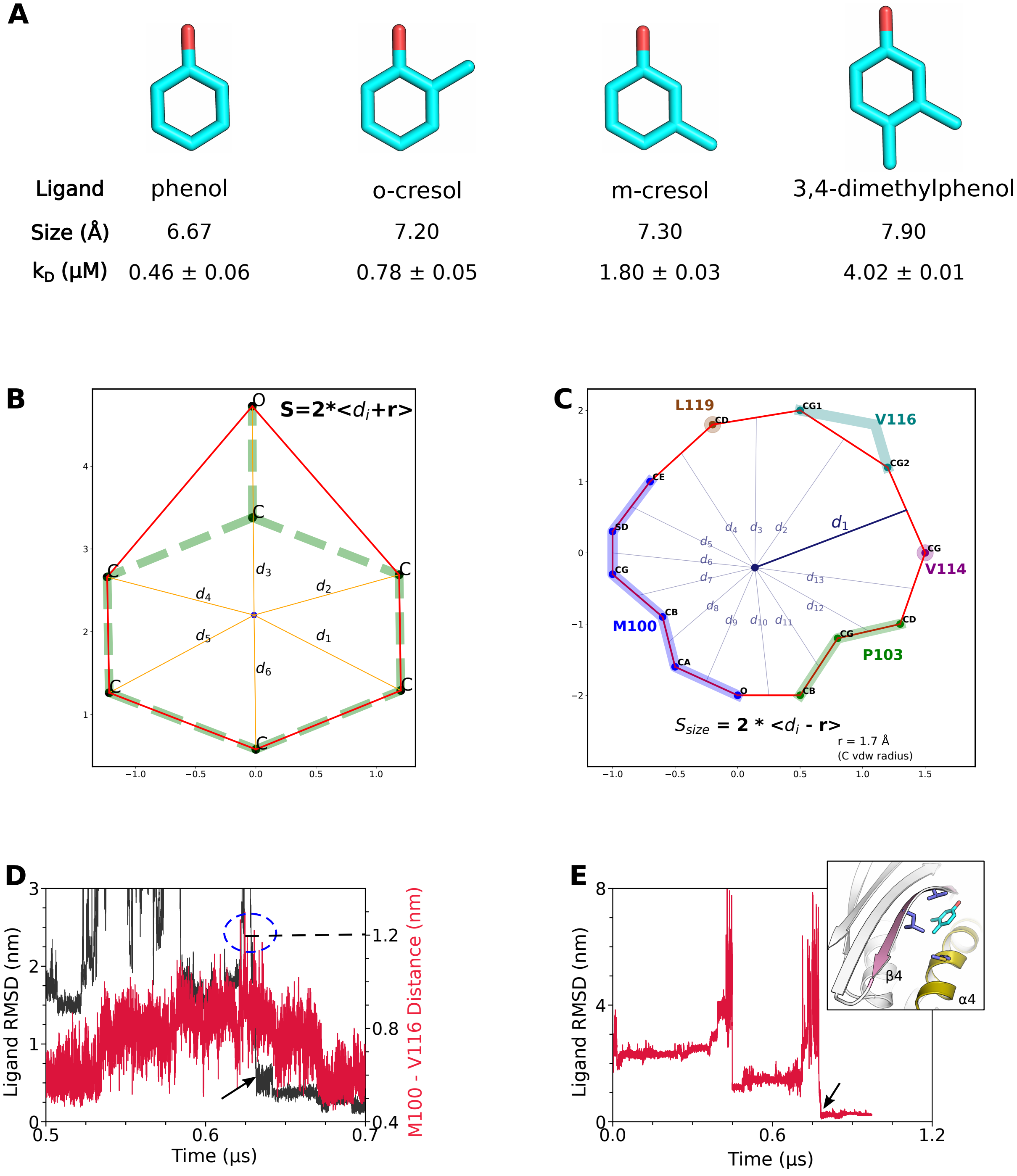


**Figure S9:** (A) Size selectivity profile of MopR with respect to different ligands. (B) **ligand size calculation**: Since aromatic ligands are planer hence to compare the ligand size with the channel opening in MopR (see maintext), the average width of ligand was used as a measure of ligand size. For each ligand's heavy atoms, the average distance from ligand's center of geometry to the outermost atoms (like convex hull) was calculated. The VDW radius of carbon atoms was also added to the distances to account for width of atoms. Twice of this average distance represents the approximate diameter of planer aromatic ligands, which was taken as their size). An example of phenol is shown. (C) **Channel size calculation**: The channel opening was ...

[... *continued S9 caption*] not perfectly circular or 2 dimensional, but we assumed it roughly circular and calculated its average diameter or channelsize. To calculate the the channel size, only the heavy atoms were considered and hydrogens were considered to be highly flexible with no role in ligand selection. The channel opening was lined by heavy atoms of M100, P103, V114, V116 and L119. Note that V114 and L119 exist in different rotameric states and anyone of their CG or CD (respectively) atom can line the channel. The heavy atoms lining the channel formed a roughly circular polygon and apothem to each side was calculated from center of geometry of channel . Van der Waals radius of carbon (1.7 $\AA$) was subtracted from apothem length to account for unavailable space due to hard spherical atoms. Twice of average apothem length was used as a measure of channelsize which can be directly compared with ligand size. Note that channel size and M100-V116 distance were perfectly correlated (linear correlation - 0.95) validating the channel size calculation. (D) M100-L119 distance duirng ligand enrty (around 1.2 nm) suitable for phenol. (E) 3,4-dimethyl phenol (3,4-dmp) binding through pathway-1. (inset) 3,4-dmp binding through pathway-1.

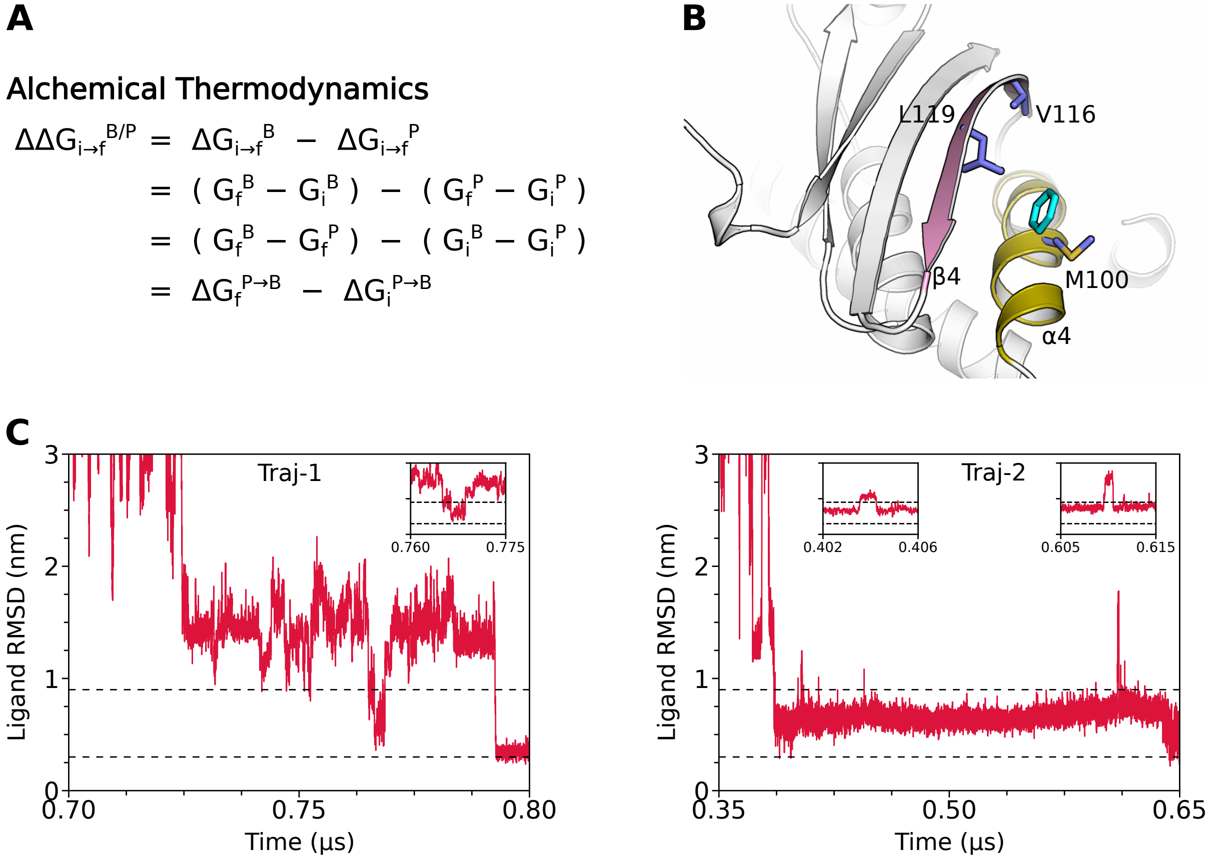


**Figure S10:** (A) Calculation of relative transition affinity of B (benzene) with respect to P (phenol) for transition from state 'i' to 'f' based on Alchemical Thermodynamic cycle. (B) Benzene binding through pathway-1. (C,D) Benzene RMSD time profiles showing benzene binding through pathway-1 and unsuccessful binding attempts (insets), with ligand (benzene) bouncing back from intermediate-2 to unbound state.

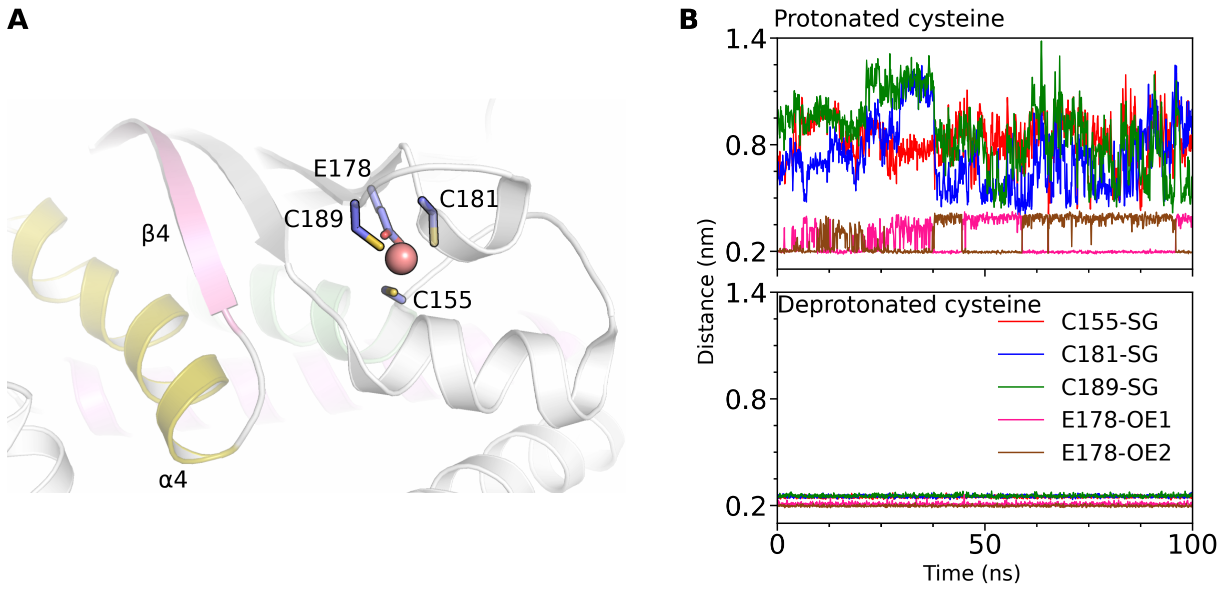


**Figure S11:** (A) $ZN^{2+}$ ion tetrahedrally coordinated to C155, E178, C181 and C189 forming $[Zn(Cys)_4]^{2-}$ complex. (B) Zinc conformation with protonated and deprotonated cystienes as measured by bond lengths between zinc and coordinating atoms. Zinc with deprotonated cysteines showed stable conformation.
